# MIND-Map; A Comprehensive Toolbox for Estimating Brain Dynamic States

**DOI:** 10.1101/2024.12.13.628351

**Authors:** Mohammadreza Khodaei, Paul J. Laurienti, Sean L. Simpson, Heather M. Shappell

## Abstract

Studying dynamic brain states has offered new insights into understanding functional connectivity. One of the promising approaches for estimating these brain states is the hidden semi-Markov model (HSMM). However, despite its potential, its adoption in the neuroscience community has been limited due to its complexity. We developed the Markov Inference Dynamic Mapping (MIND-Map) toolbox to overcome this limitation. This interactive user-friendly toolbox leverages HSMM to identify brain states, analyze their dynamics, and perform two-sample statistical comparisons of network dynamics. Furthermore, it introduces a new approach, not yet used in conjunction with these models, for determining the optimal number of states, addressing a key challenge in the field. We assessed the performance of the HSMM and our method for identifying the optimal number of states using two datasets, including a unique dataset explicitly developed for this purpose.

## 1 Introduction

The study of whole-brain functional networks, estimated from functional magnetic resonance imaging (fMRI) data, has experienced rapid growth over the past two decades, as evidenced by efforts such as the Human Connectome and the 1000 Functional Connectomes Projects [1, 2]. These studies have revealed information about the functional connections between brain regions, providing insight into the organization of brain communication in health and disease. Knowing how regions interact is vital to understanding how the brain carries out tasks involved in behavior and cognition and studying how networks differ in brain disorders is essential to identifying biomarkers and treatments. In static connectivity analyses, one (aggregate) brain network that does not change during a specific experimental scanning session is estimated. While convenient, this oversimplifies the functional relationship between brain regions. Empirical studies indicate that brain networks are dynamic and changing even in short time periods [3, 4]. Thus, dynamic brain network analyses have been introduced and provide a series of networks spanning the entire duration of a scan [5–7]. Given that the brain is a complex, dynamic system [8, 9] rather than a stationary one, dynamic brain network analyses can provide more profound insight into behavioral shifts and adaptive processes in normal and abnormal brain function.

Despite its recent popularity, dynamic brain network analysis is still very much in its infancy. The sliding-window correlation analysis is the dominant approach for generating dynamic brain network states [10]. In this technique, correlation matrices are successively computed from segments of the entire time series data, followed by their clustering into distinct states via a k-means approach [3]. However, this approach exhibits several drawbacks. First, the need to configure multiple parameters, such as window function, length, and overlap levels, poses a challenge, as suitable settings are elusive due to the absence of ground truth in fMRI data. In addition, this approach has difficulty estimating abrupt changes in connectivity. Some of the above limitations have led to the development of a new method where hidden semi-Markov models (HSMMs) were used for dynamic functional connectivity analyses [11]. The framework is ideal given that the model may be fit on a large population of subjects, is likelihood-based (which lends itself to model selection techniques), is able to estimate abrupt changes in network states, allocates a state to each timepoint, and does not assume geometric sojourn times (the continuous time a subject remains in a state before transitioning to another one).

Although the HSMM has demonstrated its effectiveness in the study of dynamic brain connectivity, its complex setup has limited its widespread adoption within the neuroscience community. Additionally, the requirement to specify the number of states prior to fitting the model poses further challenges. The number of states one fits to the data heavily influences the results of the analysis and given that there is not a ‘true’ number of brain states in real data, deciding on the optimal number of states is a difficult problem. To overcome these issues, we have developed a user-friendly toolbox (Figure 1) based on the HSMM called MIND-Map (Markov Inference Network Dynamic Mapping). This toolbox not only simplifies the use of the HSMM but also incorporates a method for choosing the optimal number of states to fit the data. The toolbox effectively captures various characteristics of dynamic connectivity, including brain states, state transition patterns, state sojourn time distributions, subject state sequences, and subjects’ state occupancy time.

**Figure 1.**
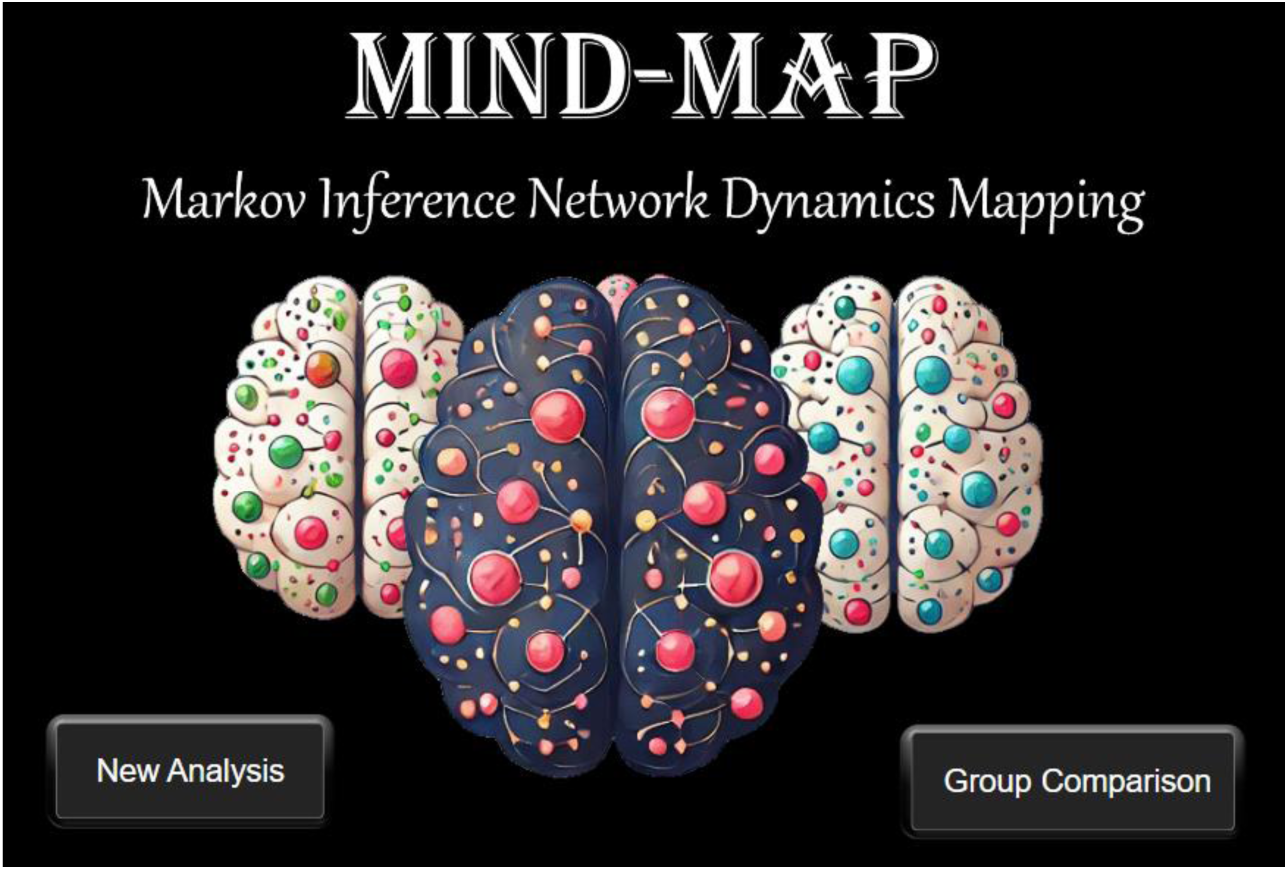
MIND-Map (Markov Inference Network Dynamic Mapping) toolbox home page.

In this article, we begin by explaining the HSMM, followed by our approach to identify the optimal number of states to fit the data. We then present a simulation study used to assess the effects of random noise on the model, something that has not yet been investigated. Moreover, our simulation study explores the impact of fitting too few or too many states and evaluates the effectiveness of our proposed method in accurately identifying the optimal number of states in the simulated data. Finally, we introduce the MIND-Map toolbox, which incorporates the HSMM analyses along with the approach for choosing the optimal number of states to fit.

## 2 The Hidden Semi-Markov Model

The standard hidden Markov model (HMM) framework assumes that an individual *i*’s time series data across all brain regions of interest during each time point *t* of an fMRI scan, denoted as *Y*_*it*_, can be described via a series of repeating brain states. Each state is characterized by its connectivity structure and its mean activation at each brain region. The collection of *p* (one for each brain region) vectors of observed time-series data for participant *i* is denoted by 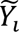. A key assumption of the HMM model is that *Y*_*it*_ comes from a multivariate normal distribution, dependent on the latent (i.e., unknown) state at time t. In other words, 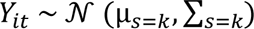, where *s*_*t*_, s ∈ {1, …, k}, denotes an index variable representing a hidden state. The vector of true network state variables for participant *i* is denoted by 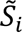,. The main goal of the HMM in this setting is to estimate a) the latent states that give rise to the fMRI data we are observing, and b) the network dynamics of how these states evolve.

The HSMM is an extension of the HMM (Figure 2). The primary distinction between the two models lies in how they handle the sojourn time distributions, the duration of time spent continuously in a state (Figure 2. Dashed lines). HMMs assume that the sojourn time distribution for each state is geometric. This suggests that the probability of remaining in a state decreases as more consecutive time is spent in a state. This results in a higher weight for shorter sojourn times in each state, which is not usually a suitable assumption for brain states. HSMMs address this issue by permitting each state sojourn distribution to be directly modeled and estimated. This includes parameterizing the sojourn distribution using models such as the Poisson distribution, gamma distribution, non-parametric distribution, or other suitable distributions.

**Figure 2.**
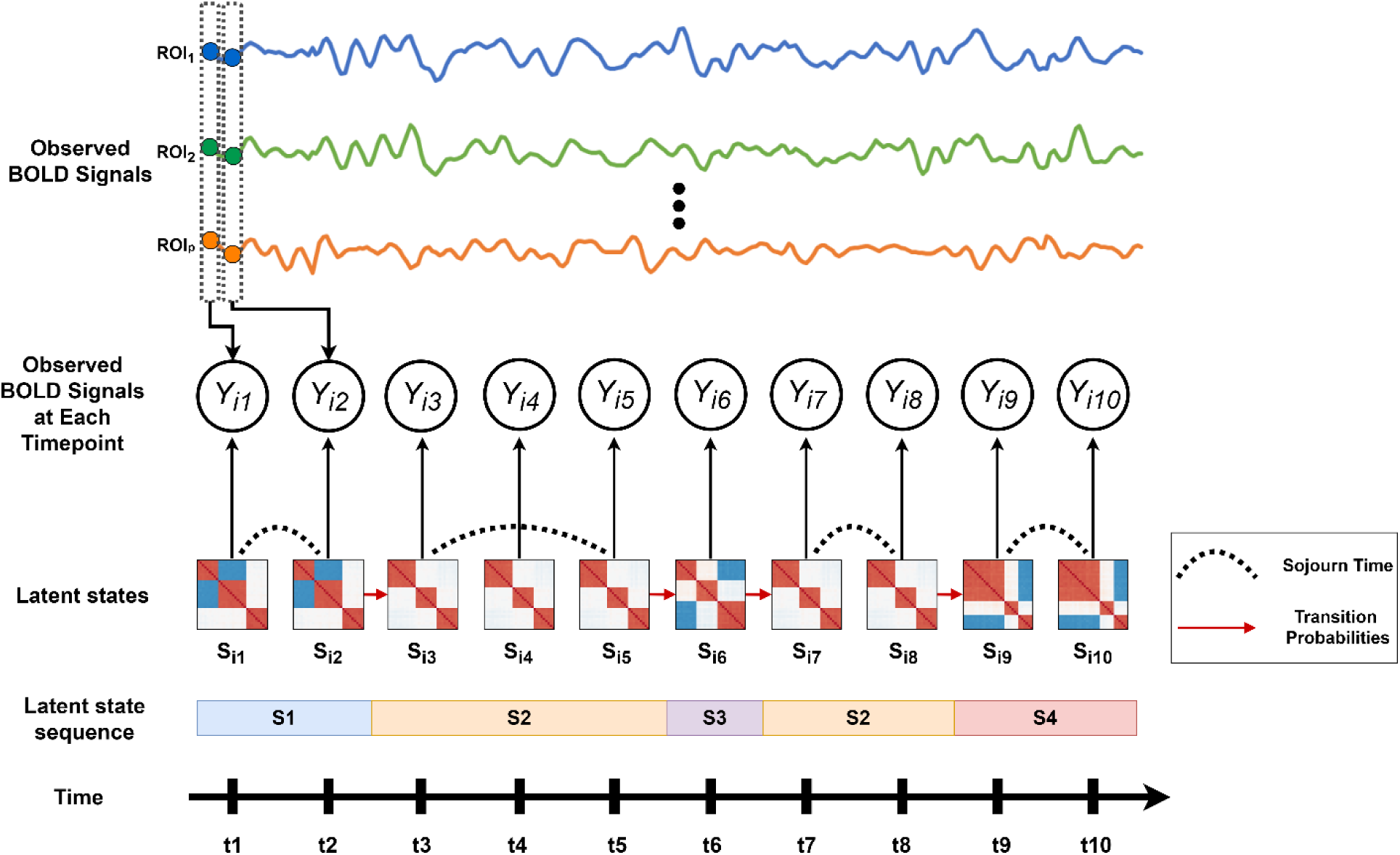
fMRI time series of p regions (*ROI*_1_, *ROI*_2_, …, *ROI*_*p*_) are illustrated. At each time point, the fMRI time series (observations), *Y*_*it*_, are associated with a corresponding hidden state covariance/correlation matrix (colored matrices). These matrices each reflect an underlying latent brain network that gives rise to the fMRI time series data we are observing. Sojourn times are depicted with dashed lines, while conditional transition probabilities are indicated by red arrows. The associated state sequence is presented at the bottom.

The HSMM takes each participant’s fMRI time series data as input and utilizes maximum likelihood estimation to calculate the states’ covariance matrices (Σ_1:*k*_), mean vectors (µ_1:*k*_), sojourn time distributions (*d*_1:*k*_(*u*)), the conditional transition probability matrix (P) (i.e., the probabilities of going to each other state when leaving the current state), and the initial state probabilities μ_1:*k*_ (the probability of starting a state sequence in a particular state). The maximum likelihood estimation routine maximizes the complete data log likelihood, which, for a single participant, can be written as follows:

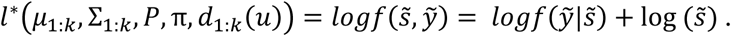

The logarithm of the above equation can be decomposed into four parts: one to model the first state, one to model the evolution of the intermediary states, one to model the final state, and one to model the conditional distribution of the observed fMRI data given the underlying latent state. In other words,

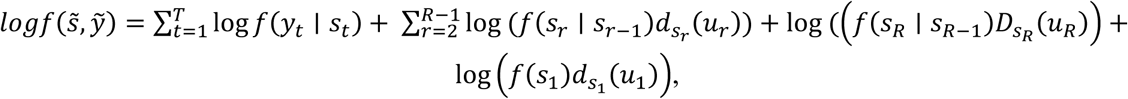

where *u*_*r*_ is the number of consecutive time points in the state. Note that

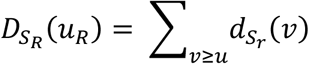

is the survivor function and models the sojourn time in the final state. It allows one to not make the assumption that the participant is leaving the final state after time *T*.

Once the model parameters are estimated, each participant’s most probable sequence of true network states may be estimated using the Viterbi Algorithm [12]. The Viterbi algorithm takes as input the state estimates, transition probability estimates, sojourn density estimates, and initial state probability estimates derived from fitting the Hidden Semi-Markov Model (HSMM) on all participants. It then computes the most probable sequence of states for each individual participant based on these estimates and their own fMRI time-series data.

We used the *mhsmm* R package [13] to estimate the model parameters and perform the Viterbi algorithm. More details about the HSMM and maximum likelihood estimation are available in Shappell et al. [11].

## 3 Selecting the Number of States

One of the primary challenges with the HSMM is the requirement to specify the number of states prior to fitting the model. An inaccurate specification of the number of states can lead to the identification of spurious or merged states. The elbow method is commonly used to identify the optimal number of states when applying a sliding window approach combined with k-means clustering to estimate hidden states [3]. However, for HMMs, there is no consensus on a standard method. Although leave-one-out Bayesian Information Criterion (BIC) [11] and Akaike Information Criterion (AIC) [14] computations have been used for HMMs, they present certain challenges. Leave-one-out BIC is time-intensive, as its typically require running the model multiple times within a leave-one-subject-out cross validation framework, making it difficult to implement for large datasets. For example, with a sample size of 100 participants, the HSMM model must be run 100 times, leaving out one participant in each iteration. On the other hand, AIC is sensitive to sample size and may become unreliable when applied to small datasets. Here we incorporated an approach for estimating the number of states grounded in our simulation study (refer to the “4. Simulation Section”). Our solution to this problem involves running the model with different numbers of states and identifying the run with the highest distance between its states. The working hypothesis behind this idea is that genuine brain states are distinct from one another, whereas the structure of spurious states resembles that of genuine brain states. In other words, the real brain states have large distances from each other, while spurious states have relatively short distances from other states. Therefore, to identify the run with no spurious states, we execute the model with varying numbers of states and subsequently assess the minimum distance between the identified states in each run. The run exhibiting the <u>highest minimum distance</u> between its states helps us pinpoint the optimal number of states, as it indicates the highest distinctiveness in the estimated states (refer to 5.3 for more details). We refer to this method as PeakMin, short for “Peak of the Minimums.”

## 4 Simulations

### 4.1 Simulation 1 (SIM-1) Data Generation

A common approach for simulating distinct brain states is generating states with unique modular structure (Figure 3, top). We used the SimTB toolbox [15] to generate 50 dynamic brain state data sets each consisting of 50 simulated subjects and five unique modular states embedded in the time series. Each subject’s simulated data included 360 timepoints and 30 regions. We selected 30 regions, rather than the default ten provided by the SimTB toolbox, to simulate data that is more comparable in size that that used in real studies. Consequently, we increased the number of time points to 360 (equivalent to a 12-minute scan with TR = 2s) to support the estimation of larger state correlation matrices. We generated a set of ‘true’ network states (correlation matrices) by randomly assigning nodes to three distinct modules in each state. We then generated a set of time series associated with each state and subsequently concatenated them to form subject time series based on a random subject sequence (Figure 3, bottom). The random sequence was generated by assigning each subject six time slots, with a uniformly distributed random numbers (from 1 to 5) assigned to each slot. These time series were convolved with hemodynamic response function (HRF), and 10% random noise was added. Random noise was generated by sampling from a standard normal distribution.

**Figure 3.**
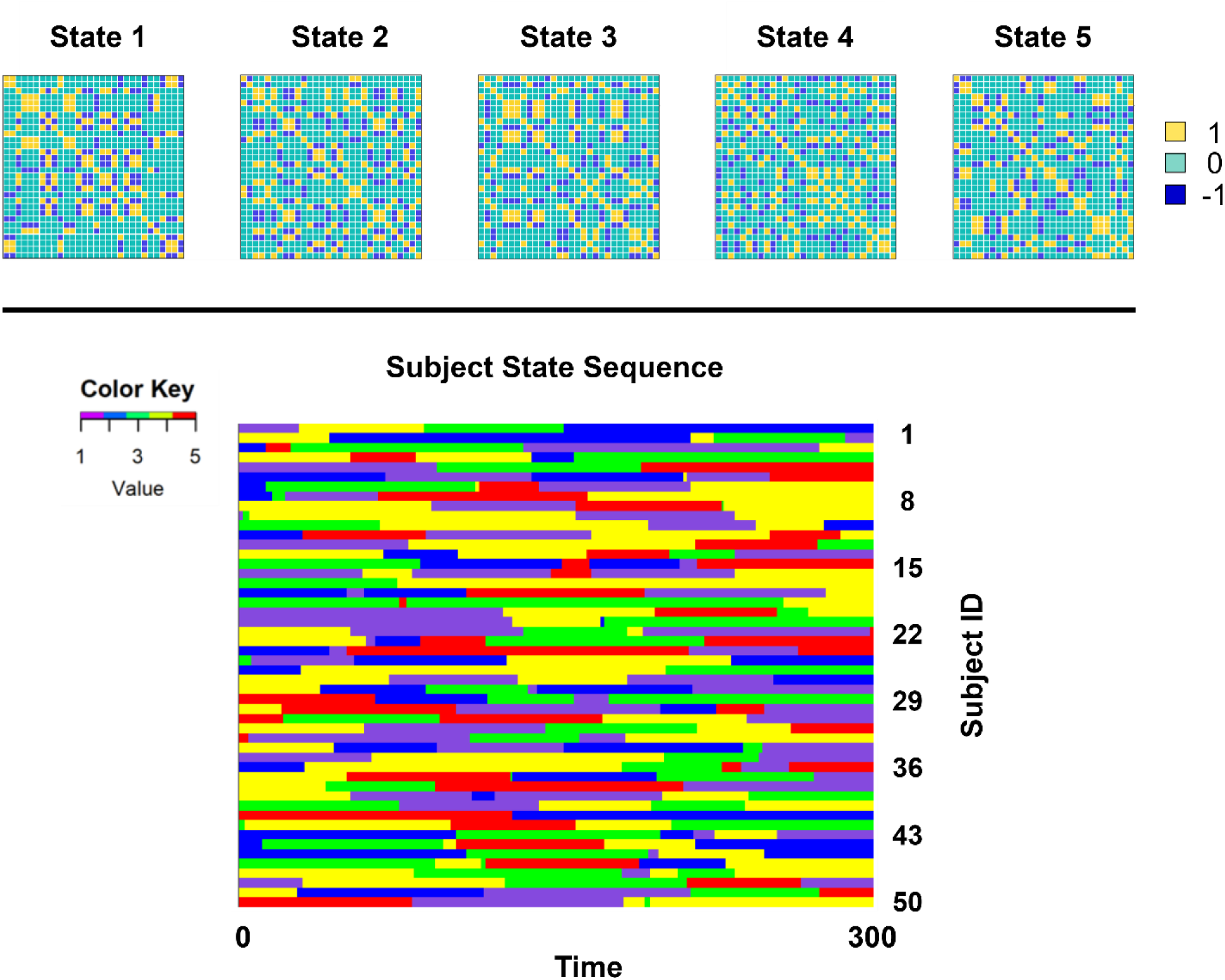
SIM-1 simulation. We generated 50 sets of groups of 50 simulate subjects’ data. Top. Illustration of one of the simulated state sets embedded in the timeseries. Bottom. Subject state sequence.

Using the approach described above, we generated 50 groups of datasets by incorporating distinct state modular structures and introducing varying random noise into the time series of each group. Throughout the remainder of this manuscript, we refer to this simulated dataset as SIM-1.

### 4.2 Simulation 2 (SIM-2) Data Generation

Although simulation using SimTB toolbox is very helpful for the initial testing of the model and simplifies the visual matching of the estimated states with the true states, the structure of these states may not accurately reflect realistic brain states. In addition, instead of random state sequences, we wanted a more realistic subject state sequence. To overcome these limitations, we generated a second set of simulated data. Instead of a highly modular state structure, we used real static brain networks as our “true” states.

To generate the new simulated data set, we began by employing five brain network correlation matrices (Figure 4. Top) estimated from the real fMRI data of 5 subjects, each containing 200 time points and 58 nodes. Each matrix was generated from a single person’s preprocessed fMRI data by performing a cross correlation on the entire time series data. The resulting five networks were used as the ground truth for each of five states. We performed a Cholesky decomposition on each state’s correlation matrix. Then multiplied the outcome by a set of independent random time series to generate a set of timeseries associated with each state [16]. However, this method is not perfectly accurate, and depending on the random time series, the resulting time series may have a correlation matrix that deviates slightly from the original. To address this, we performed multiple iterations. We performed 20,000 iterations to identify the optimal sets of independent time series that, when multiplied by states Cholesky decomposition, produce dependent time series with the least difference in correlation compared to the original correlation matrices (representing the true states). This process resulted in a collection of time series data with 200 timepoints that when processed with a cross correlation yielded networks that corresponded to each “true” state (*A*_*n*_).

**Figure 4.**
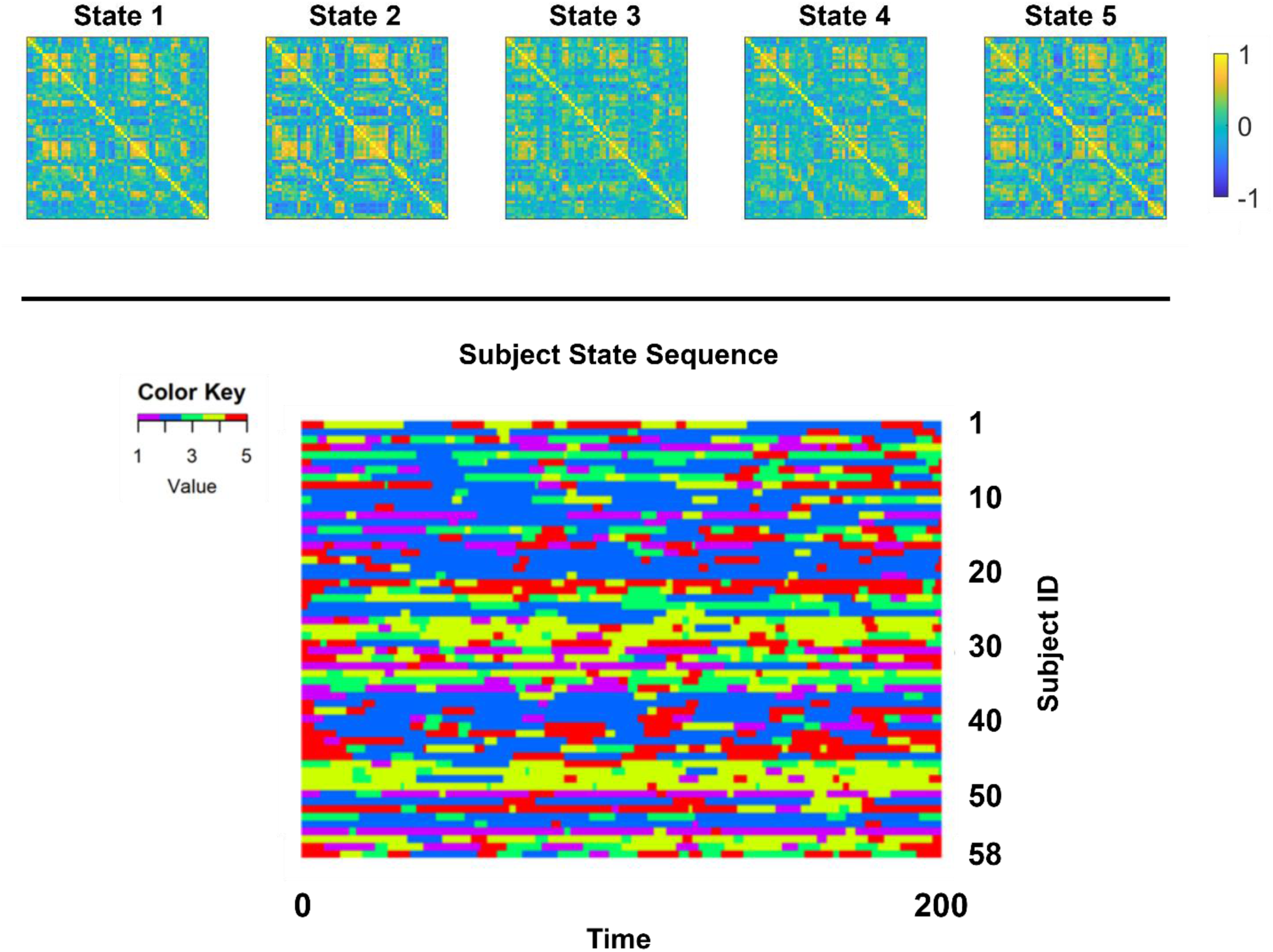
SIM-2 simulation. Top. Five real brain static network which we embedded in the simulated timeseries. Bottom. Real brain subject state sequences used to concatenate the data.

Next, we utilized the subject state sequence (Figure 4. bottom) extracted from one of our ongoing fMRI studies investigating the effects of successful weight loss in older adults [17]. Using these subject-state sequences, we concatenated different segments of the time series associated with each state to construct the subjects’ time series. We then convolved the subject time series with the hemodynamic response function (HRF) to generate the simulated fMRI signal. Lastly, N percentages of random noise, ranging from 10% to 80%, was added to (100%-N) percentage of the time series. This was achieved by generating noise with the same mean and standard deviation as the time series and then adding it to the signal at specified percentages. For instance, for 10% noise, 10% of the noise amplitude was added to 90% of the time series amplitude. We repeated this procedure 50 times, resulting in a sample of 50 sets of groups, each containing 58 subjects for each noise level. Each subject had 58 regions with 200 time points. The selection of the number of subjects, regions, and time points was also inspired by the weight loss study [17], as we aimed to investigate the model’s stability with small, real-world datasets. We will refer to this simulated dataset as SIM-2 throughout the remainder of the manuscript. Figure 5 demonstrates the pipeline for generation of SIM-2.

**Figure 5.**
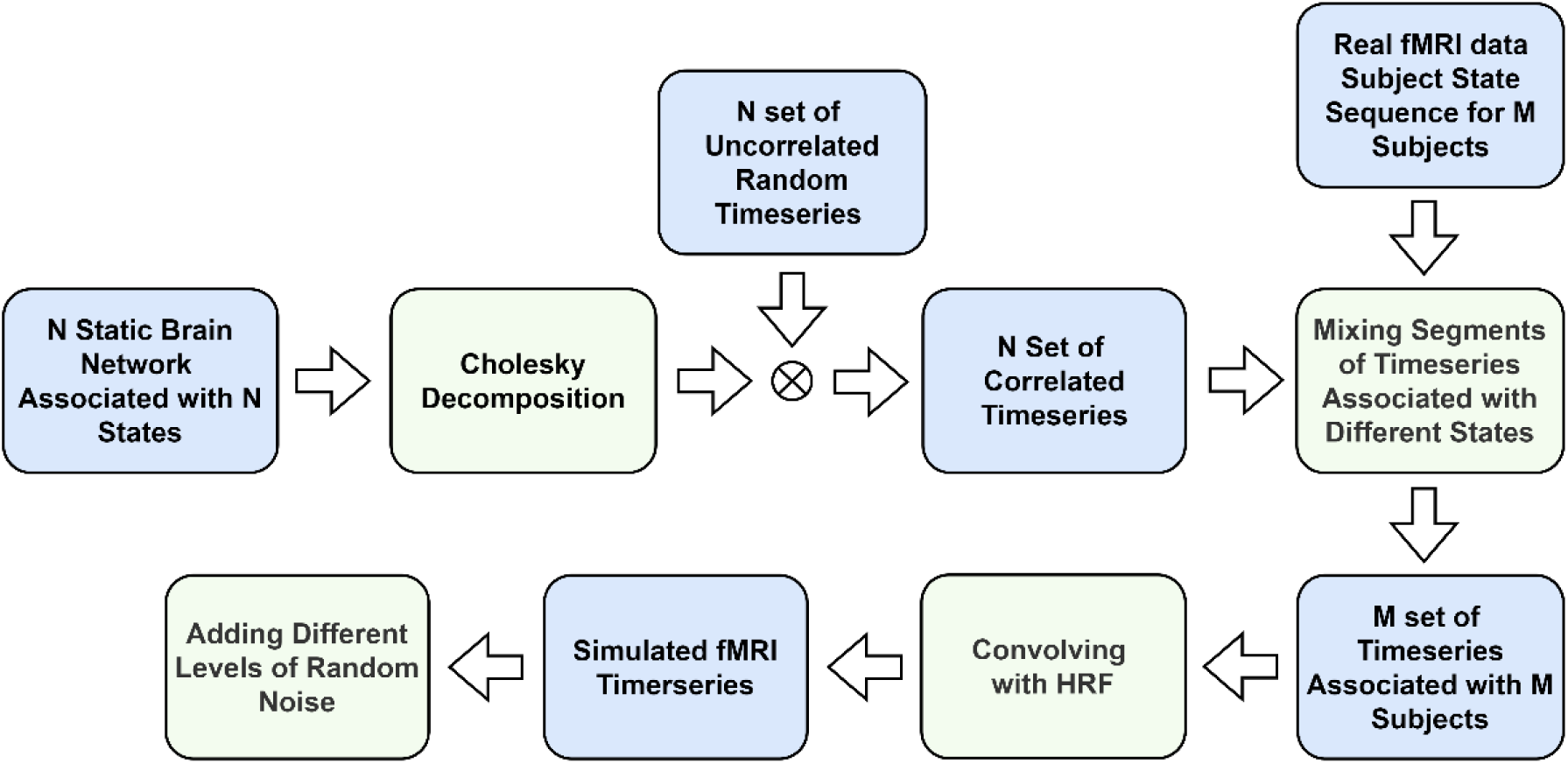
The pipeline for generating SIM-2 simulated data.

### 4.3 Assessment of the HSMM using Simulated Data

To evaluate the performance of the HSMM, we focused primarily on SIM-2, as we believed it presented a greater challenge to the model compared to SIM-1. We used SIM-2 datasets to examine the HSMM’s effectiveness in identifying the true subject state sequence under two conditions: (1) varying levels of noise and (2) fitting an incorrect number of states. To assess our approach’s accuracy in identifying the true number of states, we analyzed both SIM-1 and SIM-2 datasets. Each HSMM analysis applied the k-smoothed nonparametric distribution for sojourn time modeling. Figure 6 provides details for each simulated dataset and explains how they were utilized.

**Figure 6.**
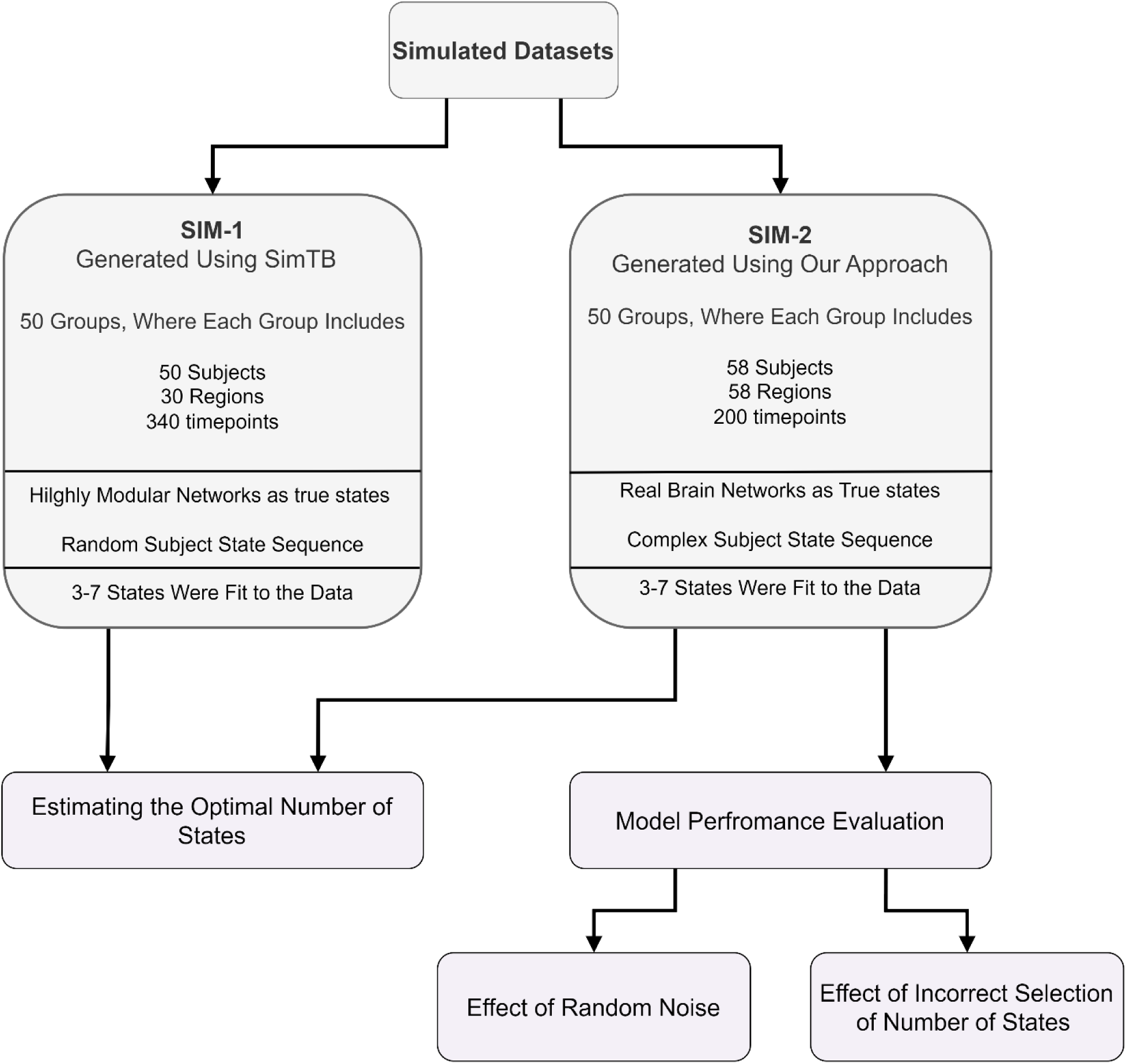
We used two simulated datasets, SIM-1 and SIM-2, in our simulations to evaluate our approach for identifying the true number of states. SIM-2, being a more realistic model of brain dynamics, was also used to assess the model’s performance under varying levels of noise and incorrect state selection. The details of each simulated dataset are provided.

#### Effect of Random Noise

We examined the model performance under varying noise levels, ranging from 10% to 80%, on the SIM-2 dataset. The simulated timeseries were fed to HSMM to estimate the states and subject state sequence. We investigated the overlap between the actual and the estimated subject state sequences across 50 runs of each noise level. Specifically, we calculated the percentage of time points where the true and estimated state sequences were in the same state. The matching was performed simultaneously for all states. In HSMM, states labels are arbitrary, meaning that two identical runs can yield different state labels (e.g., one run may label states as 1, 2, 3, 4, 5, while another may label them as 5, 2, 4, 3, 1). To identify matching states, we compared the true subject state sequence with all possible label permutations and identified the permutation with the highest state sequence match.

#### Effect of Incorrect Selection of the Number of States

In addition to examining the effect of noise, we investigated the impact of applying the HSMM with an incorrect number of states. We ran the HSMM on the SIM-2 dataset with a range of states, from 3 to 7, to evaluate its performance when an incorrect number of states was chosen, compared to when the correct number was selected. This selection was made because two states were considered too simplistic and using more than seven did not produce stable estimates due to our small datasets. This analysis was conducted on sample with 10% noise.

#### Performance of Our Approach in Estimating the True Number of States

Finally, we used both SIM-1 and SIM-2 to evaluate the performance of our approach in identifying the true number of states. SIM-1 consisted of 50 groups, where each group consisted of 50 subjects. The true number of states was 5 in all groups. For each group, we applied HSMM with varying numbers of states, ranging from 3 to 7. We then computed the distance between the state correlation matrices and counted how often the run with the correct number of states (5) had the highest minimum distance between its states. The same analysis was performed on SIM-2, which also included 50 groups, each also having five true states. We computed the distance between the states using Euclidean distance for both data sets. We also applied other distance measures, including distance measures based on the Manhattan distance, belief propagation [18] and spectral distance [19], but the Euclidean distance performed the best, so we focus our results on ones that used Euclidean distance.

## 5 Simulation Results

To illustrate the performance of the HSMM for estimating the true state structure, we applied the model to two single groups of SIM-1 and SIM-2. Figure 7 illustrates the results of applying the HSMM with the true number of states (5 states) on a single group (from 50 generated groups) from the SIM-2 dataset. As demonstrated, the model successfully estimated the true state structures, despite minor differences caused by the added 10% noise. Similarly, applying the HSMM to a single set of the SIM-1 dataset produced comparable results (see Supplementary Figure 1).

**Figure 7.**
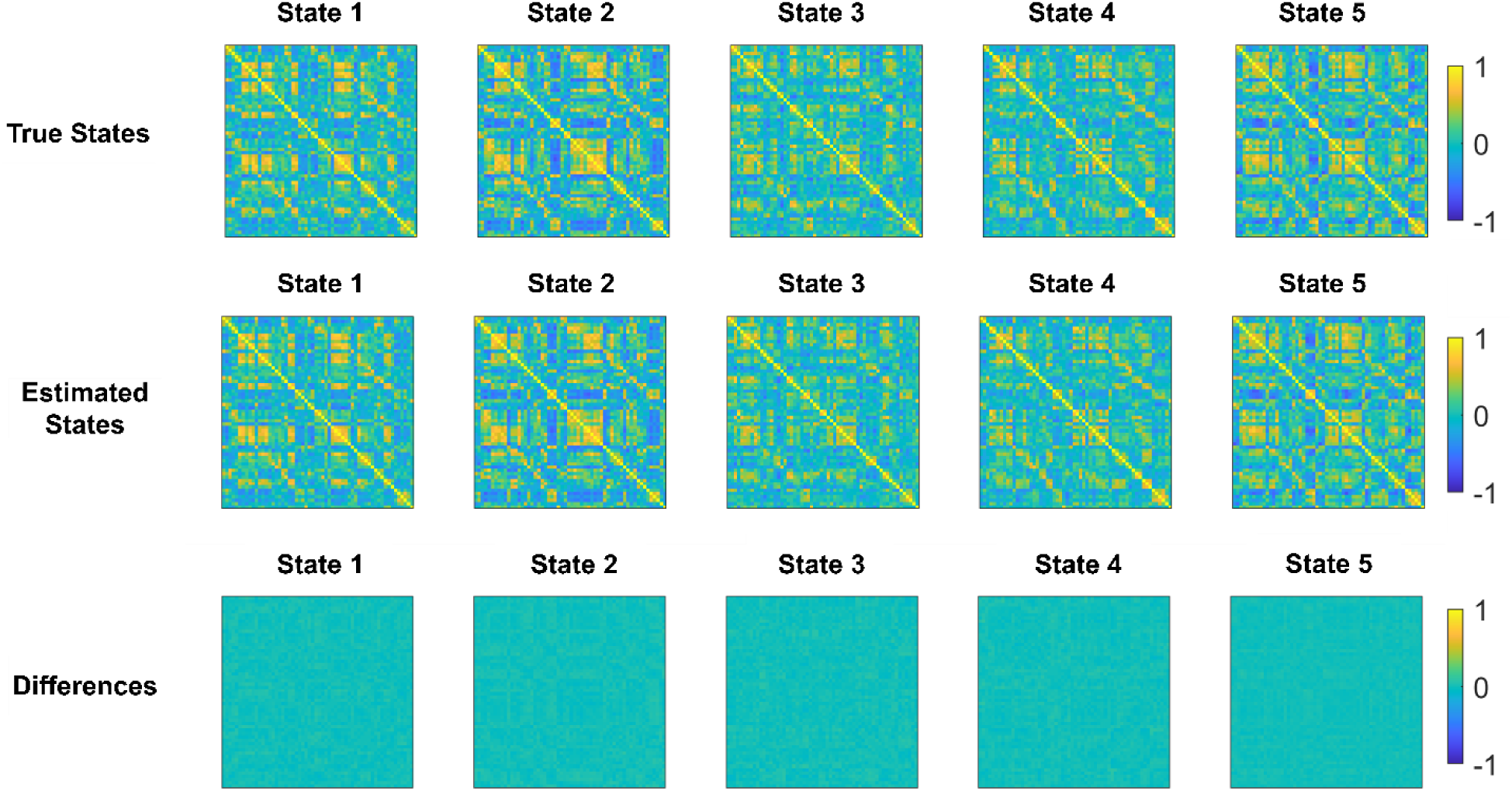
Performance of the HSMM on a single set of the SIM-2 dataset which includes 10% percent random noise. Top. True states correlation matrices. Middle. The estimated states. Bottom the difference between the true states and estimated states.

### 5.1 Effect of Random Noise on Model Estimations

We evaluated the HSMM’s performance by examining the overlap between the estimated and the true subject state sequence using the SIM-2 dataset (refer to section 4.3). First, we focused on the model’s performance in the presence of different noise levels ranging from 10% to 80%. Figure 8a illustrates the true subject state sequence alongside examples of the estimated subject state sequence (Figure 8b) for data with 20%, 40%, 50%, and 60 % noise. Figure 9c demonstrates the plot of the average overlap between the true subject state sequence and the estimated subject state sequence across 50 runs (50 groups), for each level of noise. As shown, the model accurately estimates the true subject state sequence with up to 50% noise. Beyond this point, its performance declines significantly, with 60% noise exhibiting a pattern similar to the initial state sequence setup. Figure 9 illustrates the estimated states obtained by applying the HSMM to a single group (selected from the 50 generated groups) under varying levels of noise.

**Figure 8.**
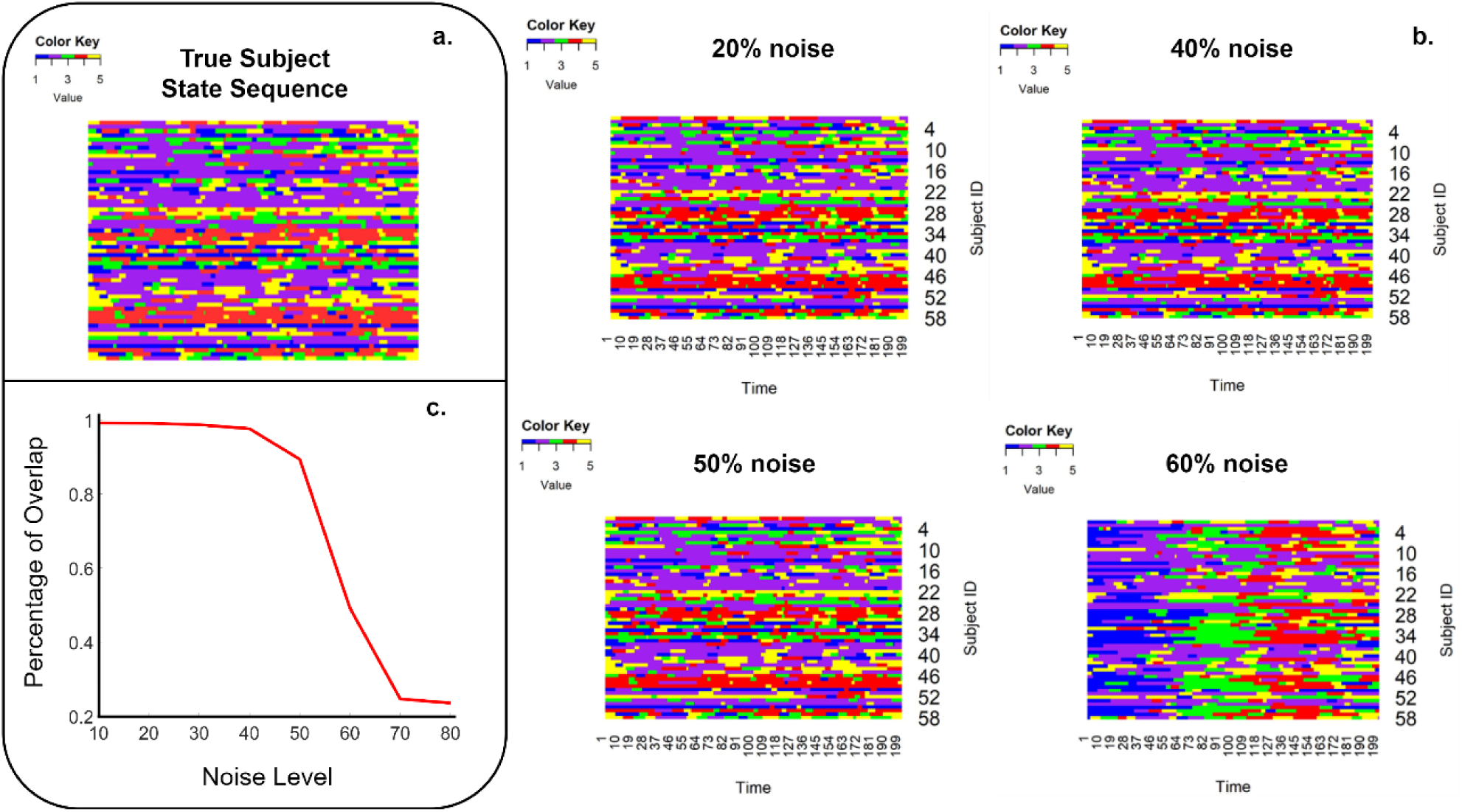
The performance of the model in the presence of random noise. a. The true subject state sequence. b. Example of estimated subject state sequences in the presence of varying noise levels. c. Average percentages of overlap between the estimated and the true subject state sequence (50 runs for each noise level).

**Figure 9.**
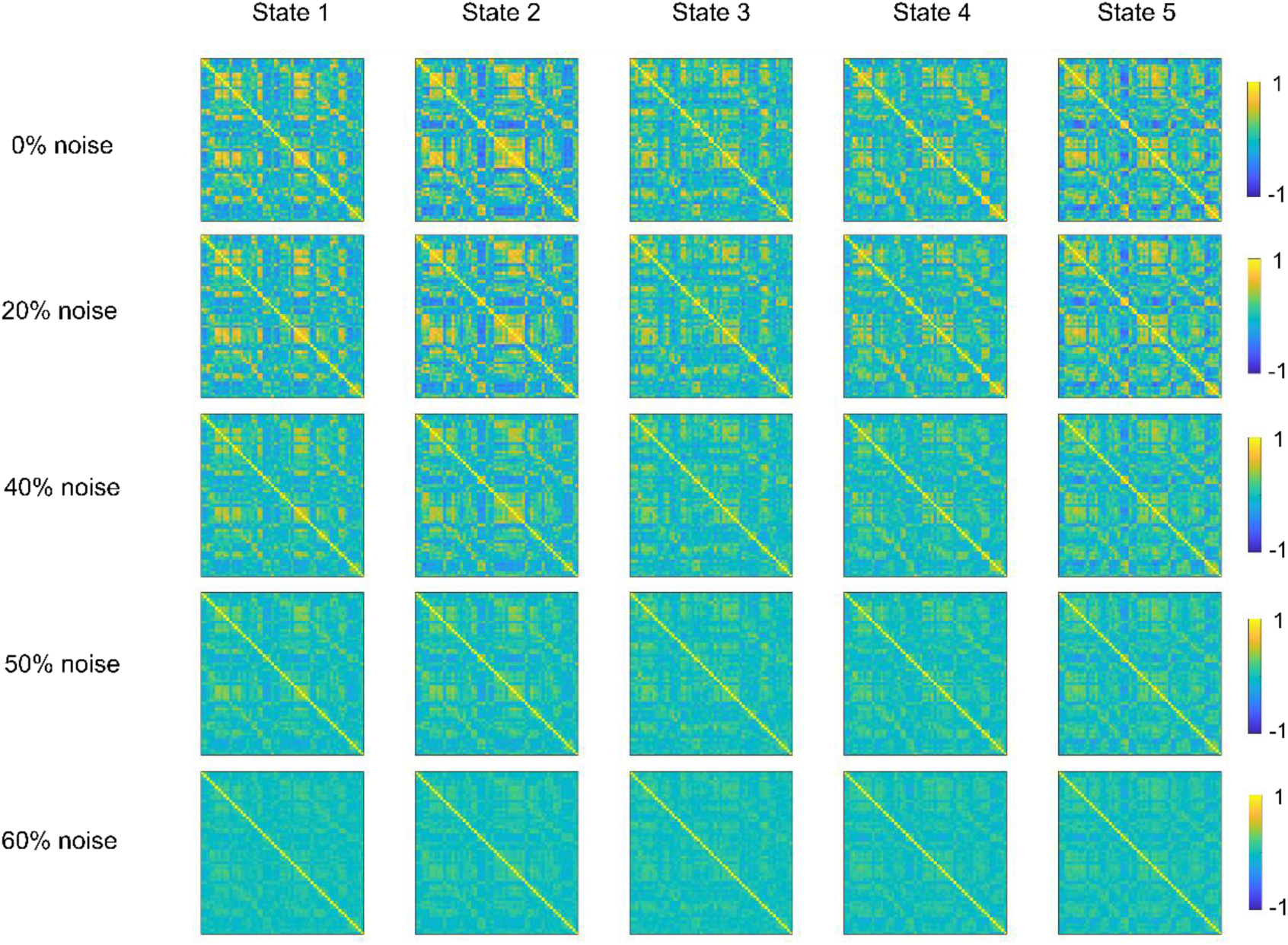
An example of the estimated states for each level of noise.

### 5.2 Effect of Incorrect Selection of the Number of States

To investigate the effect of applying the HSMM with an incorrect selection of the number of states, we conducted a HSMM analysis with varying numbers of states ranging from 3 to 7 using a simulated data with 5 states (Figure 10a). Figure 10b presents examples of subject state sequences for four runs, corresponding to runs with 4, 5, 6, and 7 states. When the HSMM runs with the true number of states (five), it performs well. It is nearly identical to the true subject state sequence. However, when it is performed with fewer states (four), the model retains some states and merges the others. As shown in the figure, states three (green) and five (yellow) are merged into one (yellow), while the other states remain largely unchanged. When more states were used (six and seven), it generates spurious states that do not occupy a large amount of time points as indicated by the black and white colors. We investigated the average percentage of overlap between the true and estimated state sequences across 50 runs (50 groups) for each number of states (refer to section 4.3). As shown in Figure 10c, the model accurately identifies the true subject state sequence only when the correct number of states (5) is used, emphasizing the importance of the number of states selection when applying the HSMM.

**Figure 10.**
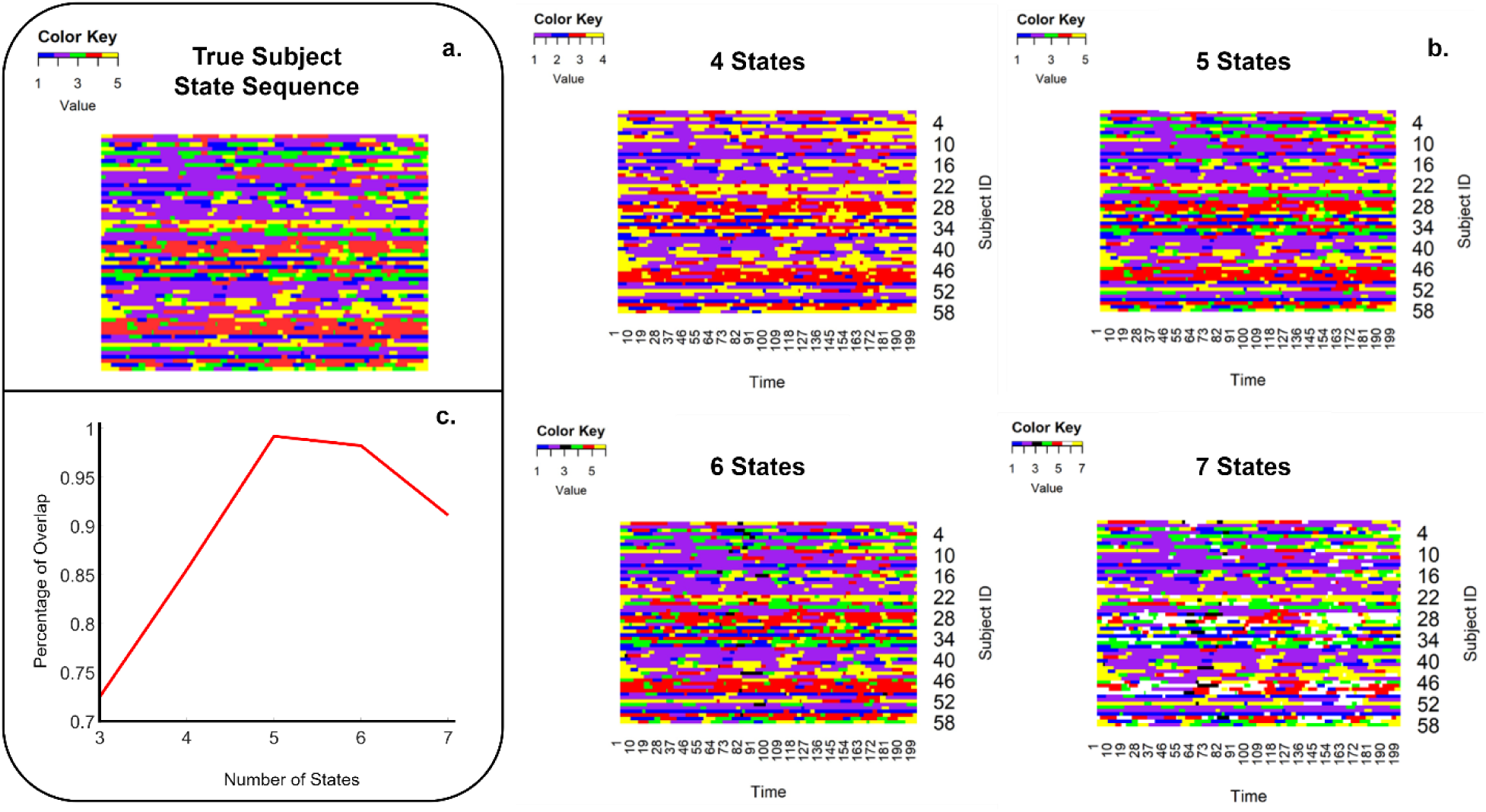
The performance of the model when HSMM is performed with different numbers of states. a. The true subject state sequence. b. Examples of estimated subject state sequences when the model is applied with different numbers of states. c. Average percentages of overlap between the estimated and the true subject state sequence (50 runs for each noise level).

### 5.3 Performance of Our Approach in Estimating the True Number of States

We propose identifying the true number of states by running the model with different numbers of states and selecting the run that yields the highest minimum distance between state pairs (the PeakMin approach). Figure 11 provides an example that illustrates why the PeakMin approach effectively determines the true number of states. In this example, the true number of states is 4, and the model is run with 3, 4, and 5 states. When the model is run with 3 states, some true states are merged, resulting in composite states with varying contributions from the original states. Conversely, when the model is run with 5 states, it estimates an extra state, using information from the true states to generate the additional state. As shown in the example, the estimated state 5 closely resembles state 3. However, when the true number of states is chosen, the model successfully identifies distinct states that differ significantly from one another. Thus, by assessing the highest minimum distance between state pairs across runs with varying numbers of states, we can reliably estimate the optimal number of states.

**Figure 11.**
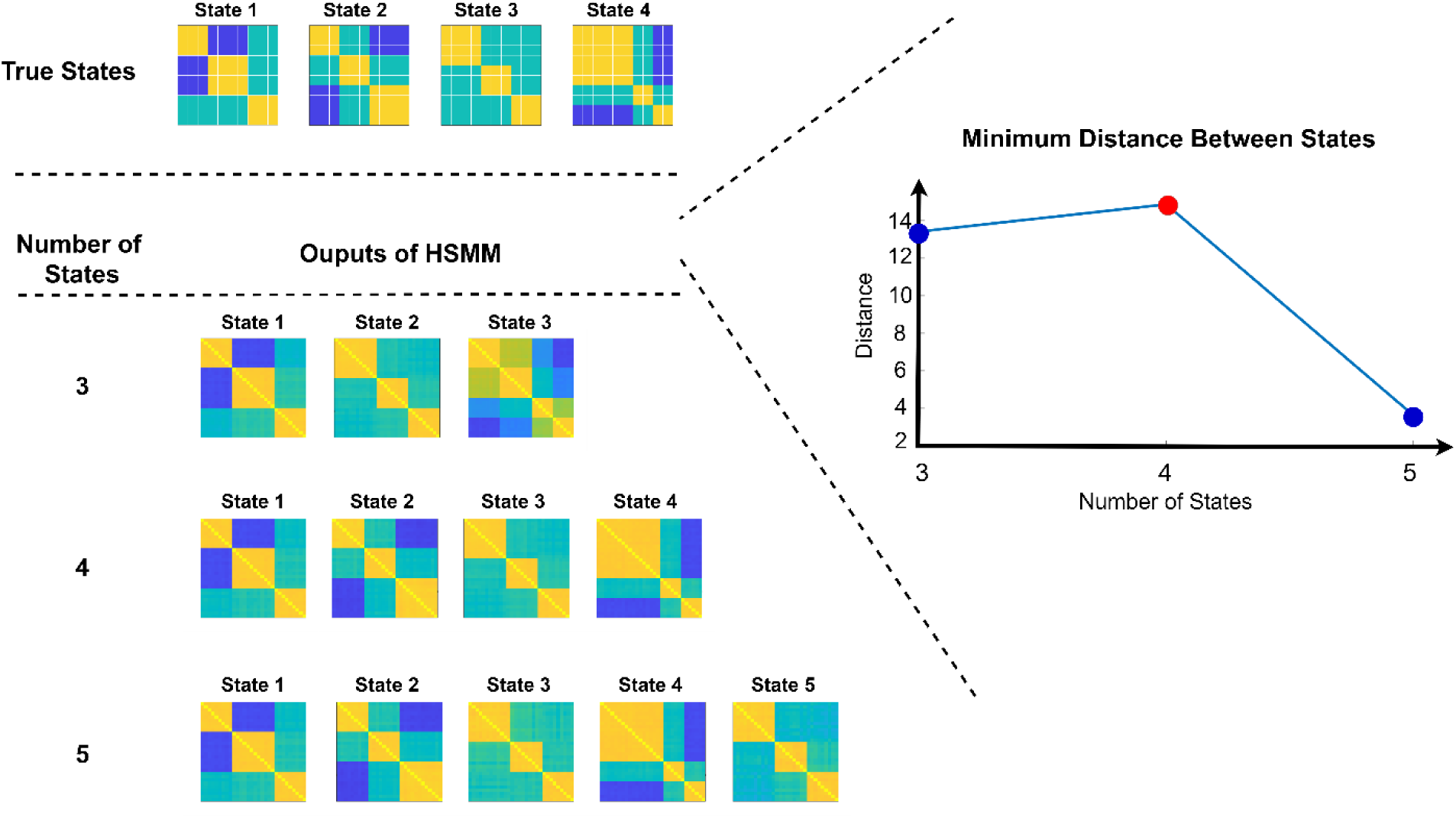
Analysis of simulated data to demonstrate our method of selecting the number of states to fit to our data. Top: The scenario where the actual number of states is four. The true four state correlation matrices are presented. Bottom: Outcomes from executing HSMM with different numbers of states (3, 4, 5). Right: Plot of minimum distance between state pairs in each run. The run with the highest minimum distance between the pairs specifies the optimal number of states.

We evaluated the performance of the PeakMin approach for estimating the true number of states using both the SIM-1 and SIM-2 datasets. A HSMM analysis was conducted with varying numbers of states for each simulated dataset. For the SIM-1 dataset, we explored a range of 3 to 7 states. Our method accurately identified the optimal number of states (5) across all successful runs as shown in Figure 12a. The plot indicates that when the HSMM is run with five states, it achieves the highest minimum distance between states compared to when it is run with other numbers of states. However, this is true only when the HSMM successfully identifies the true states (see Supplementary Figure 2 for details).

**Figure 12.**
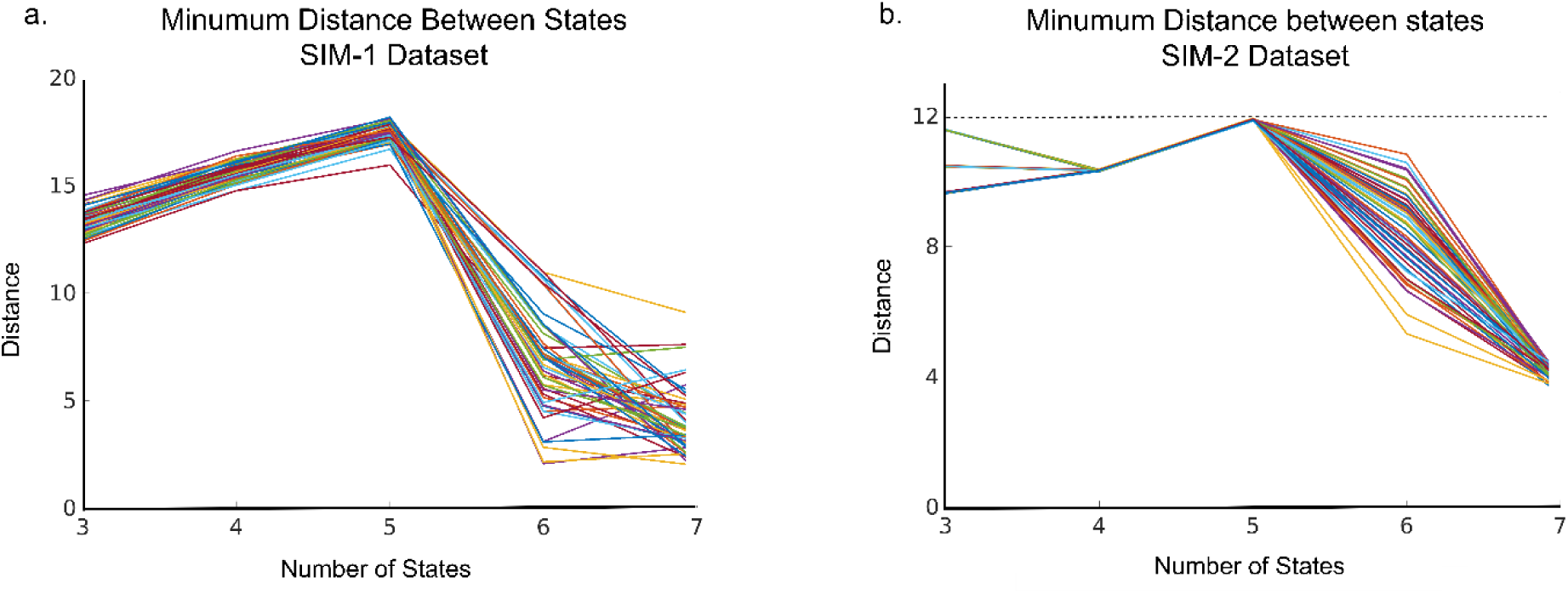
Estimating the true number of states based on the distance between the states. The run with the true number of states shows the highest distance between the states when it is compared to the others. a. Results of dataset SIM-1, which included 50 groups. In all groups, the conducted HSMM with a true number of states (5 states) had the highest minimum distance between its states. b. Results of dataset SIM-2, which also included 50 groups. In all groups, the HSMM run with the true number of states (5 states) had the highest minimum distance between its states.

For SIM-2, despite its more complex structure, the model exhibited similar performance across all runs (50 groups). The highest minimum distance between state pairs was observed when the correct number of states (5) was used in the HSMM (Figure 12b). Due to insufficient data in some runs, the model was unable to converge with more than seven states, so the analysis was limited to this range.

## 6 Mind-Map Toolbox

The pipeline of analyses implemented in the HSMM toolbox is illustrated in Figure 13. The input of the toolbox is the preprocessed fMRI time series of P brain regions. The user must either set up their preferred configuration or proceed with the default setup. This setup includes specifying the method for initializing the model parameters, sojourn time distribution type, range of number of states, and the topological measures to be estimated for each state.

**Figure 13.**
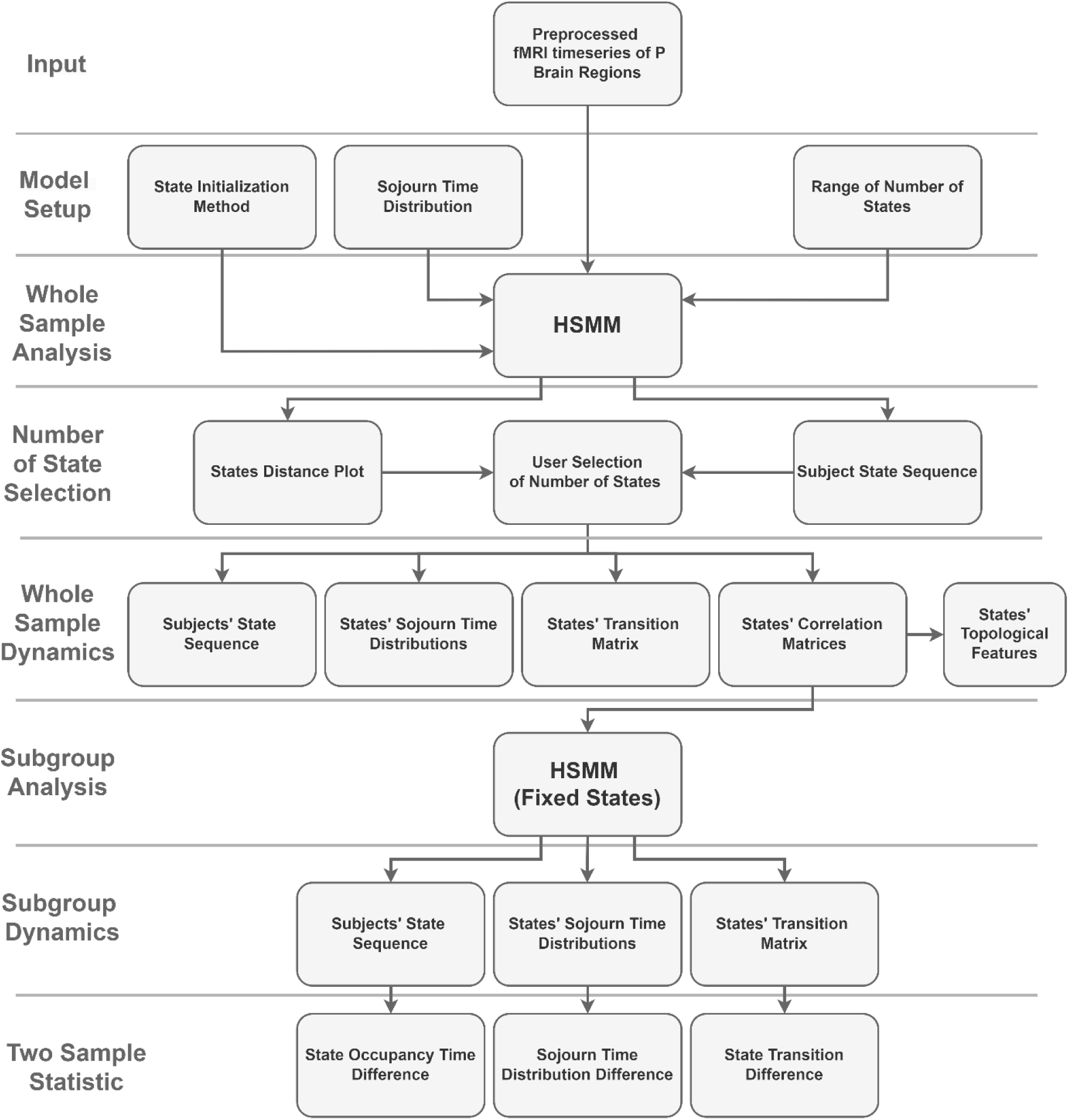
Pipeline of MIND-Map toolbox.

### 6.1 Initial Setup

The HSMM requires the initialization of several parameters, including initial state probabilities, state transition probabilities, mean vectors, covariance matrices, and sojourn distributions. In the toolbox, the state probabilities and transition probabilities are initialized with equal values for all states. Specifically, each state has an equal probability of being the starting state and equal likelihood of transitioning to any other state when a transition is occurring.

#### States Initialization Method

Initializing the states’ covariance matrices and mean vectors is more challenging, as inaccurate initialization can prevent the model from converging to the optimal states. The toolbox supports two methods for state initialization: time series equal partitioning and sliding window with k-means clustering. The time series equal partitioning approach divides each time series into k equal segments, where k is the number of states. Then, the covariance matrix and mean vector for each segment are estimated. The sliding window approach involves extracting the correlation matrix from each window, followed by k-means clustering of these matrices to identify cluster centers, which represent the states. We used tapered, overlapping sliding windows with a length of 11 time points and a shift of 1 time point. We selected window sizes that are half the length suggested in the original manuscript [3] of this approach to better capture fast transitions. While these smaller windows can be less stable, only the centers of the estimated windows are used to initialize the HSMM. Therefore, minor instabilities in the smaller windows are not expected to cause significant issues.

#### Sojourn Time Distribution

As previously mentioned, a key advantage of the HSMM is its capability to model the distribution of sojourn times. The toolbox offers three options for sojourn time distribution modeling: the nonparametric, K-smoothed-nonparametric, shifted Poisson, and gamma distributions. The nonparametric distribution models the sojourn time without the assumption of any parametric form, which enables capturing complex and unknown forms of state distribution. The K-smoothed-nonparametric is an extension of nonparametric distribution that leverage a smoothing kernel to estimate the approximate smooth distribution from the observed data. The Poisson distribution was defined with random lambda parameters between 5 and 20. For the gamma distribution, we used random values between 1 and 5 for the shape parameter and 10 and 50 for the scale parameter. Given that we often do not know the shape of our sojourn time distributions prior to fitting the model, the K-smoothed-nonparametric and nonparametric distributions are particularly useful for a first pass at running the analysis. However, compared to gamma and shifted Poisson distributions, K-smoothed-nonparametric distributions have much higher number of parameters to be estimated; therefore, more data is required. One useful strategy in the absence of enough data is to run the model with a K-smoothed-nonparametric or nonparametric distribution, identify the sojourn time distribution’s overall shape, compare its similarity to the gamma and shifted Poisson distribution, and then rerun the model with the most similar distribution.

#### Range of Number of States

The toolbox tackles the challenge of selecting the number of states by performing the HSMM with varying numbers of states and selecting the most suitable model as outlined in the section 3. The users should specify a range of the number of states to fit to their data, considering that a high number of states may not be appropriate for smaller sample sizes. The current version of the toolbox allows a range from 3 to 15 states.

#### Selection of topological measures

The toolbox estimates the network topological measures related to each state, including global efficiency [20], shortest path [21], clustering coefficient [20], local efficiency [20], assortativity [22], modularity [23, 24], strength [25], and density, using the brain connectivity toolbox (BCT) [26]. The user can select the desired topological network measure to be estimated for the states.

The HSMM analysis will run on the whole sample with the user-defined initial setup (Figure 14) using *mhsmm* R package [13] and it returns whole sample specific parameters estimates using the Expectation-Maximization algorithm [27]. Following estimation of all model parameters, the toolbox estimated each participant most probable state sequence using the Viterbi Algorithm [12]. This algorithm takes the fMRI data and all estimated model parameters including states mean vector and covariance matrices, transition probabilities, sojourn density and initial states probabilities and estimates each participant most likely state sequence. More details regarding this approach can be found in Shappell et al. [11]. Since a range of the number of states has been specified, the model will compute the dynamics for all variations of the number of states. The states’ correlation matrices and subject state sequence from each run will later be used to find the optimal number of states.

**Figure 14.**
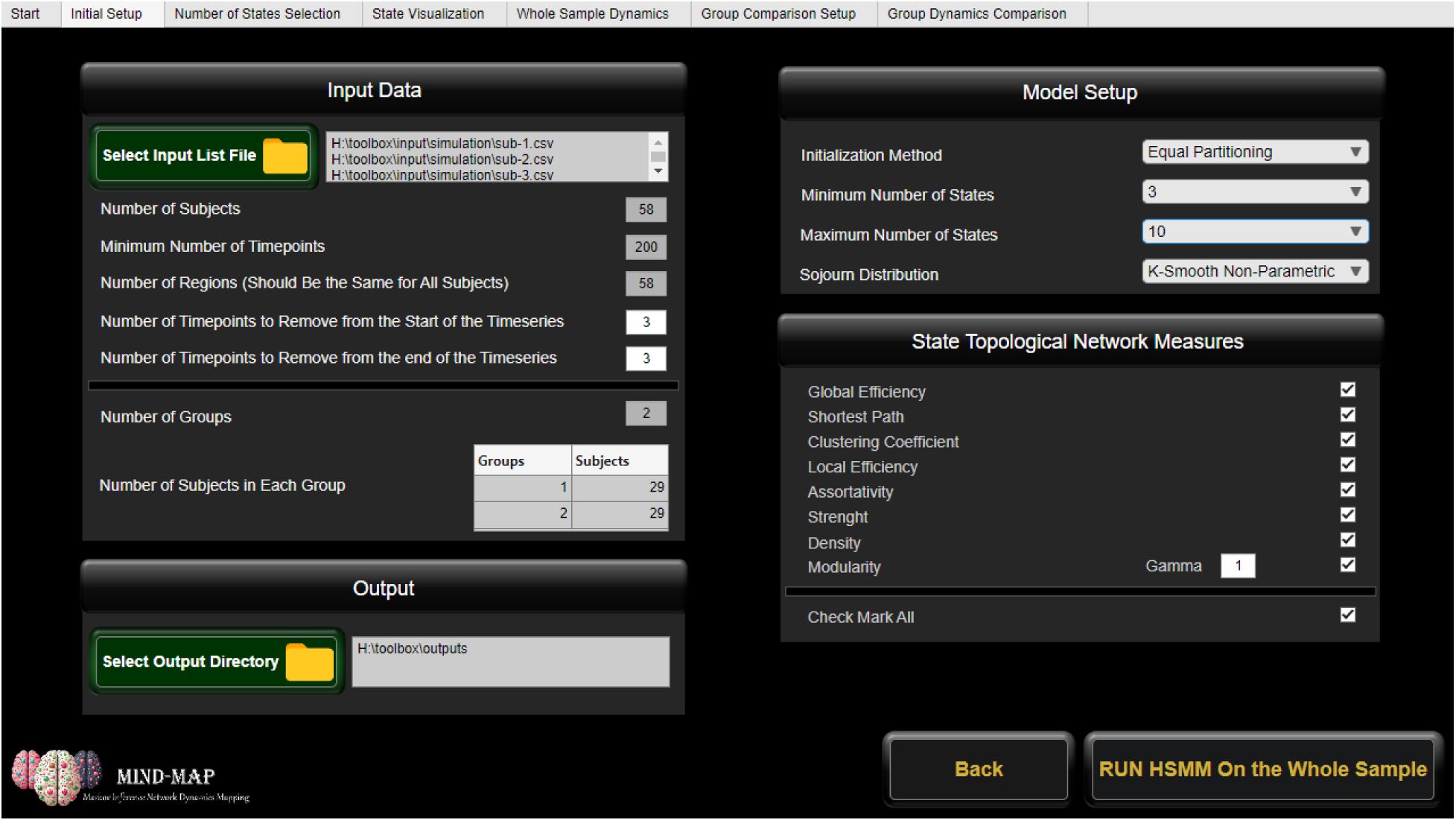
Initial Setup Tab.

### 6.2 Number of States Selection

As discussed earlier, one of the primary challenges with the HSMM is the requirement to specify the number of states. An inaccurate specification of the number of states can lead to the identification of spurious or merged states. We recommend using the PeakMin approach, which involves running the model with varying numbers of states and selecting the run that achieves the highest minimum distance between the identified states (refer to section 5.2). The toolbox computes and plots the states’ minimum distance plot, helping the user identify the most optimal number of states.

Although this approach successfully identified the correct number of states in two sets of simulated data (refer to the section 5.3), its effectiveness depends on the HSMM being appropriately fitted to the data. As outlined in the section 2, the HSMM estimates several parameters, including covariance matrices, mean vectors, sojourn time distributions, and the transition probability matrix. Additionally, involving more regions of interest (ROIs) increases the size of the covariance matrices and mean vectors, necessitating the estimation of more variables. Therefore, adequate input data (adequate sample size and length of time series) is crucial for estimating all these parameters.

Without sufficient samples, the model will fail to accurately fit the data. One output that can reflect a poor fit is the estimated subject state sequences. The subject state sequence figure illustrates the most probable state that each subject occupied at each time point of the scan. When the data is insufficient for the model to run correctly, the subject state sequence figure may illustrate a single state for most subjects throughout the whole scan time. In addition, if equal portioning has been used as the initialization method, it might reflect a state sequence close to the initial partitioning rather than the optimal dynamic state sequence. To address this issue, the subject state sequence figure is also presented in the “number of states selection” section. The user can simultaneously observe the state distance plot and the subject state sequence figure to assess whether the model accurately fits the data (Figure 15). The user can select the number of states that best fits the data based on the distance plots and subject state sequence fit.

**Figure 15.**
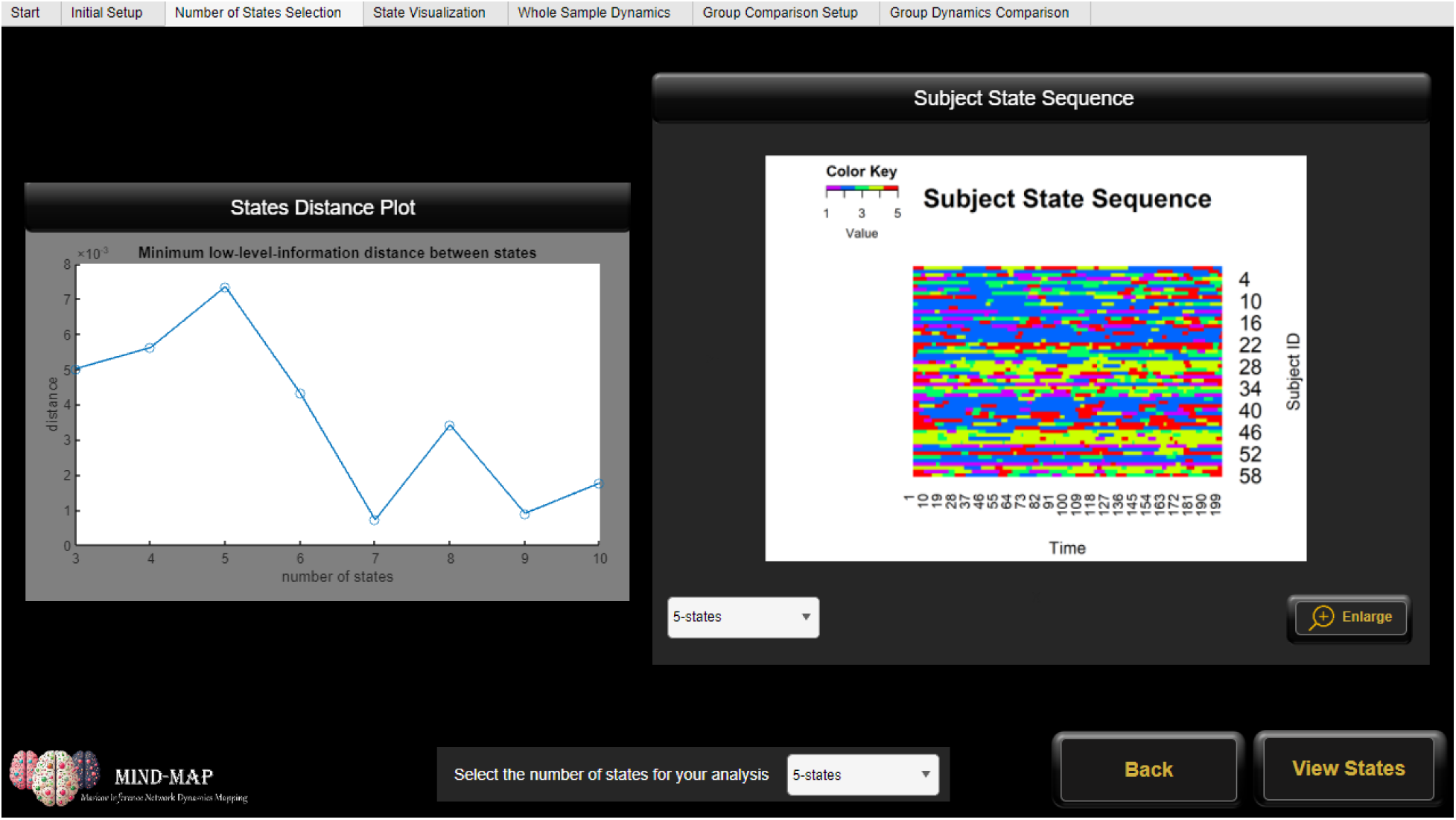
Number of States Selection tab.

### 6.3 State Visualization

Following the selection of the number of states, the states will be presented in the “State Visualization” tab (Figure 16). This tab illustrates states’ correlation matrices. In addition, the user can visualize the states on the brain using two approaches. In the first approach, the user can observe the mean activation associated with each state (Figure 17, top). The mean activation vector is one of the outputs of the HSMM model, which represents the mean z-scored fMRI signal for each ROI during each state. The mean vectors can then be mapped to the brain to illustrate the states. In addition to the mean activation visualization, the toolbox enables visualization of the modules related to each state. ROIs of each state will be divided into connected partitions using Newman’s spectral modularity [24] analysis and then mapped to the brain (Figure 17, bottom). For both brain illustrations of the states, the toolbox uses MRICrongl software [28]. The topological measures associated with each state are also present in this tab. The user can use these measures to investigate the topology differences between the states.

**Figure 16.**
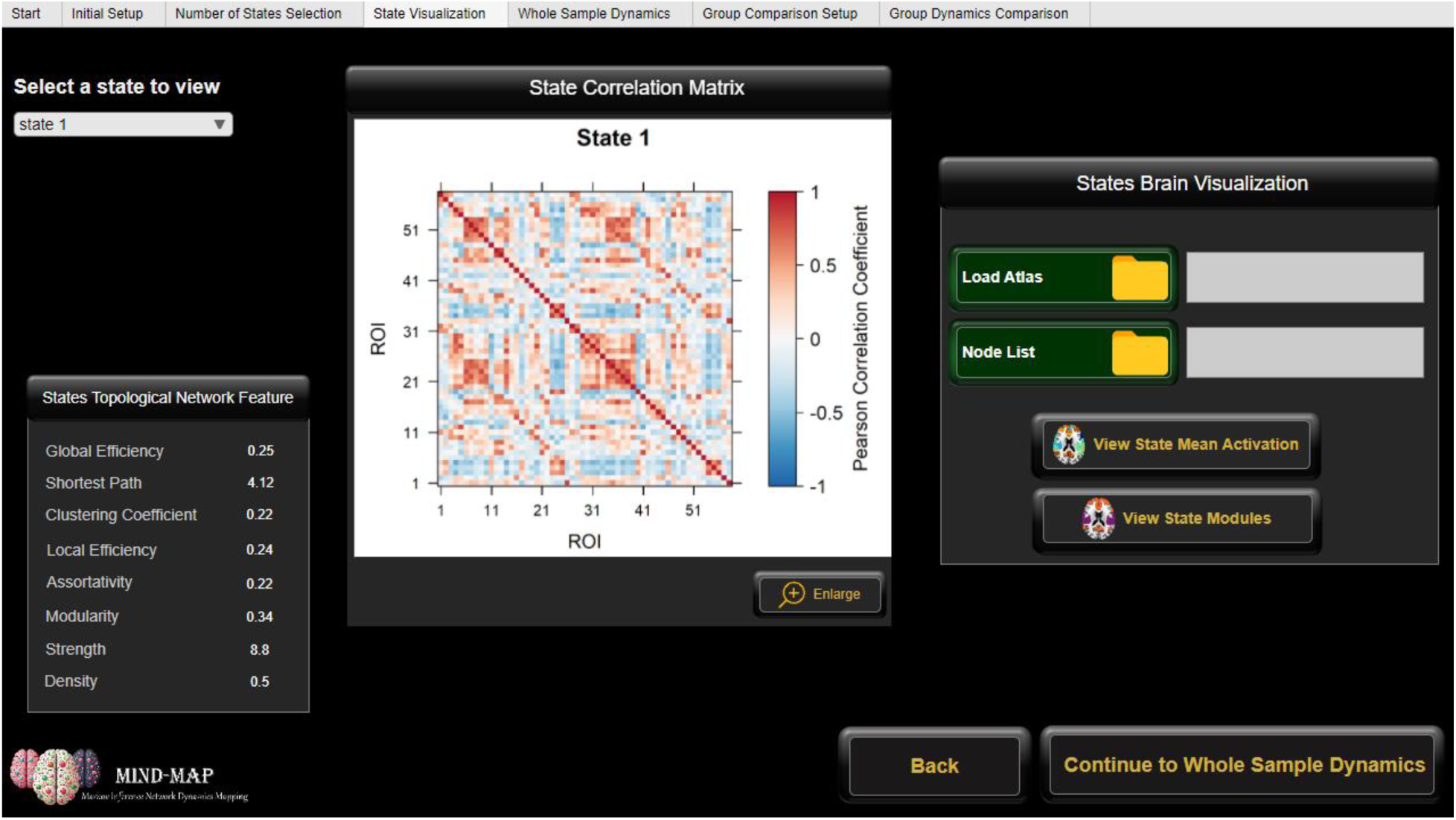
State Visualization tab.

**Figure 17.**
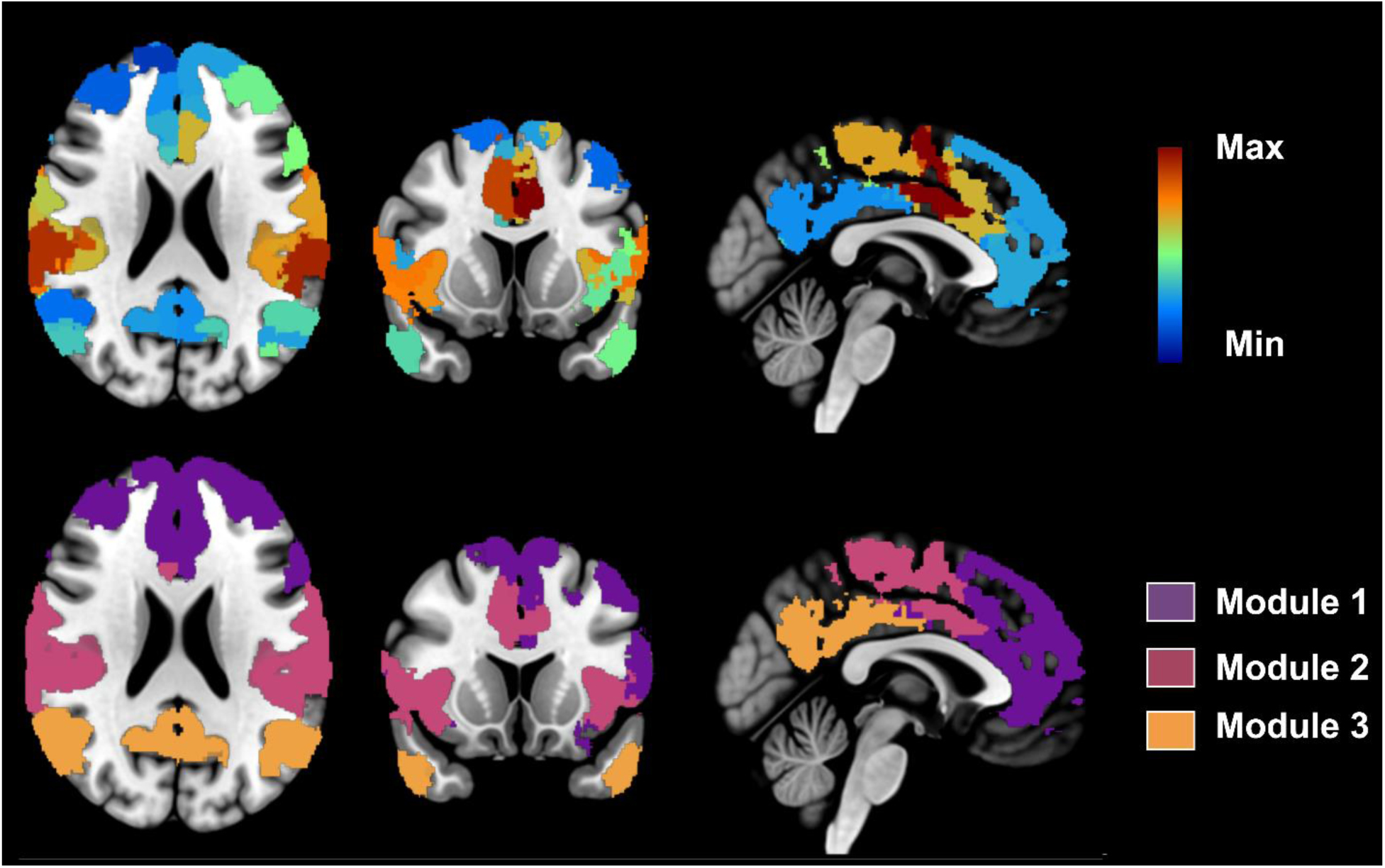
A state means activation map (top) and a state modules map (bottom) generated by the toolbox for the selected ROIs (58 out of 114) of the brain.

### 6.4 Whole Sample Dynamics

The toolbox generates a set of outputs associated with the whole sample dynamics, including sojourn time distributions for each state, mean occupancy time, state transition probabilities, and subject state sequences (Figure 18). In addition to the illustration of results, the toolbox saves the data related to each output so the user can use them to find their association with their specific covariate of interest. Each subject occupancy time and number of transitions to each state can also be found in the output directory.

**Figure 18.**
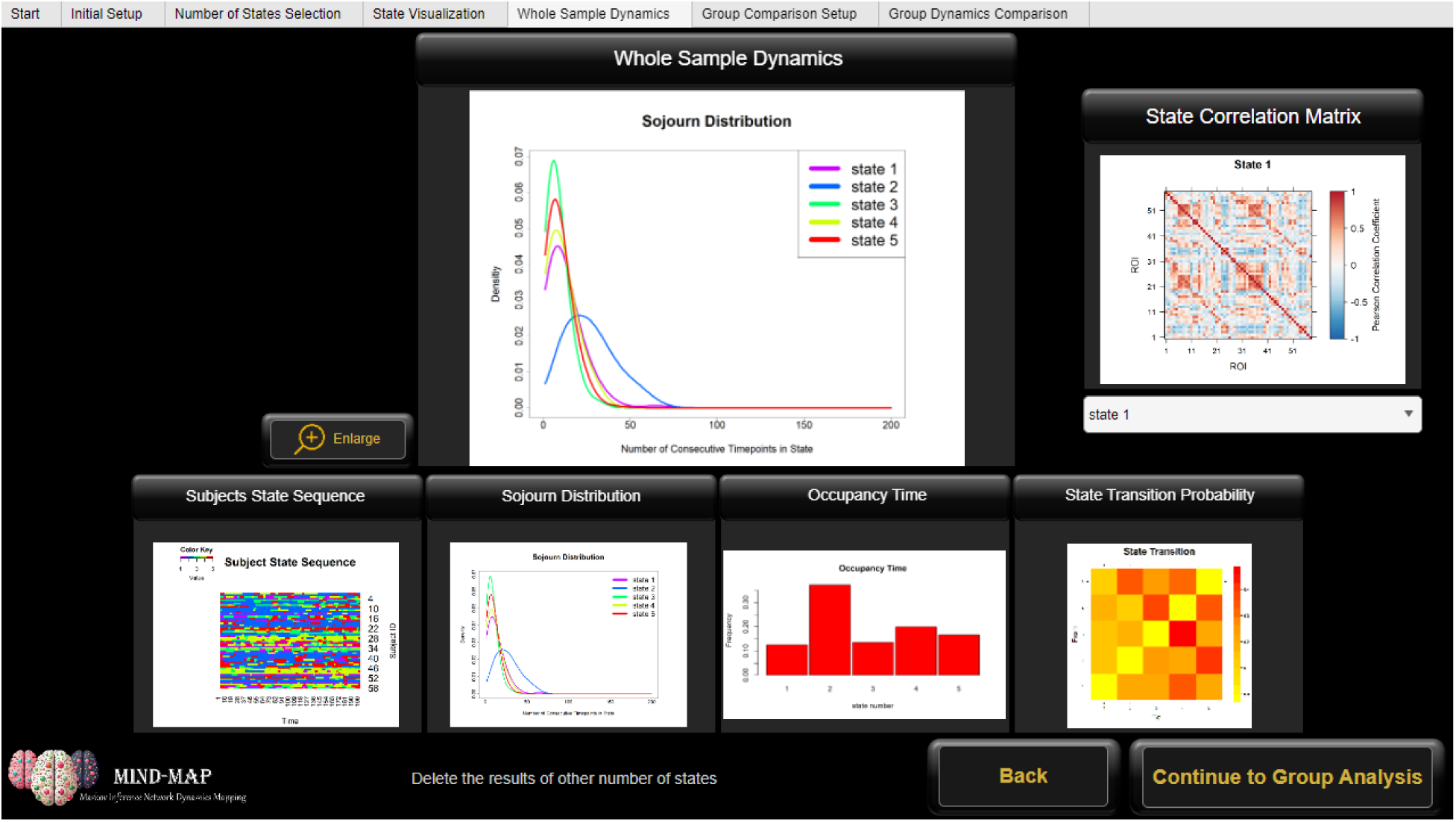
Whole Sample Dynamics. This includes sojourn time distributions, subject state sequences, state occupancy times and state transition probabilities.

### 6.5 Subgroup Dynamics

For studies with multiple groups, the toolbox allows for the extraction of the state dynamics of the subgroups by fitting the HSMM model to each group separately. For this analysis, it uses the whole sample covariance matrices and mean activation vectors (states) as input to the model. To ensure group compatibility, it fixes the states so that the model does not estimate new ones. This guarantees all groups have the same states, so their dynamics are comparable. This process generates subgroup-specific state dynamics, which are then displayed in the group comparison tab, highlighting the differences in sojourn time distributions, occupancy times, transition matrices, and subject state sequences.

### 6.6 Subgroup Comparison Statistical Testing

The current version of the toolbox allows for statistical comparisons between two groups, focusing on comparing, for each state, the empirical sojourn distribution, subject occupancy time, and subject number of transitions from and into particular states (Figure 19). The empirical sojourn distribution for each subject is estimated by counting the number of consecutive timepoints each subject spends in each state. These counts are then aggregated for each group, resulting in a distribution of consecutive time points associated with each state for each group. The empirical sojourn distributions of each state are compared between the two groups using kullback-Leibler (KL) Divergence [29], with 500 permutation. The KL test is performed using the Philentropy R package [30]. For paired permutation testing, the toolbox randomly swaps corresponding participants’ subject state sequences between the two groups, generates new empirical sojourn distributions for groups, and compares them. This process is repeated to create a distribution from all runs, which is then used to compare the actual group difference. For unpaired testing, participants are randomly permuted between the two groups, and then the same process is followed as in paired permutation testing.

**Figure 19.**
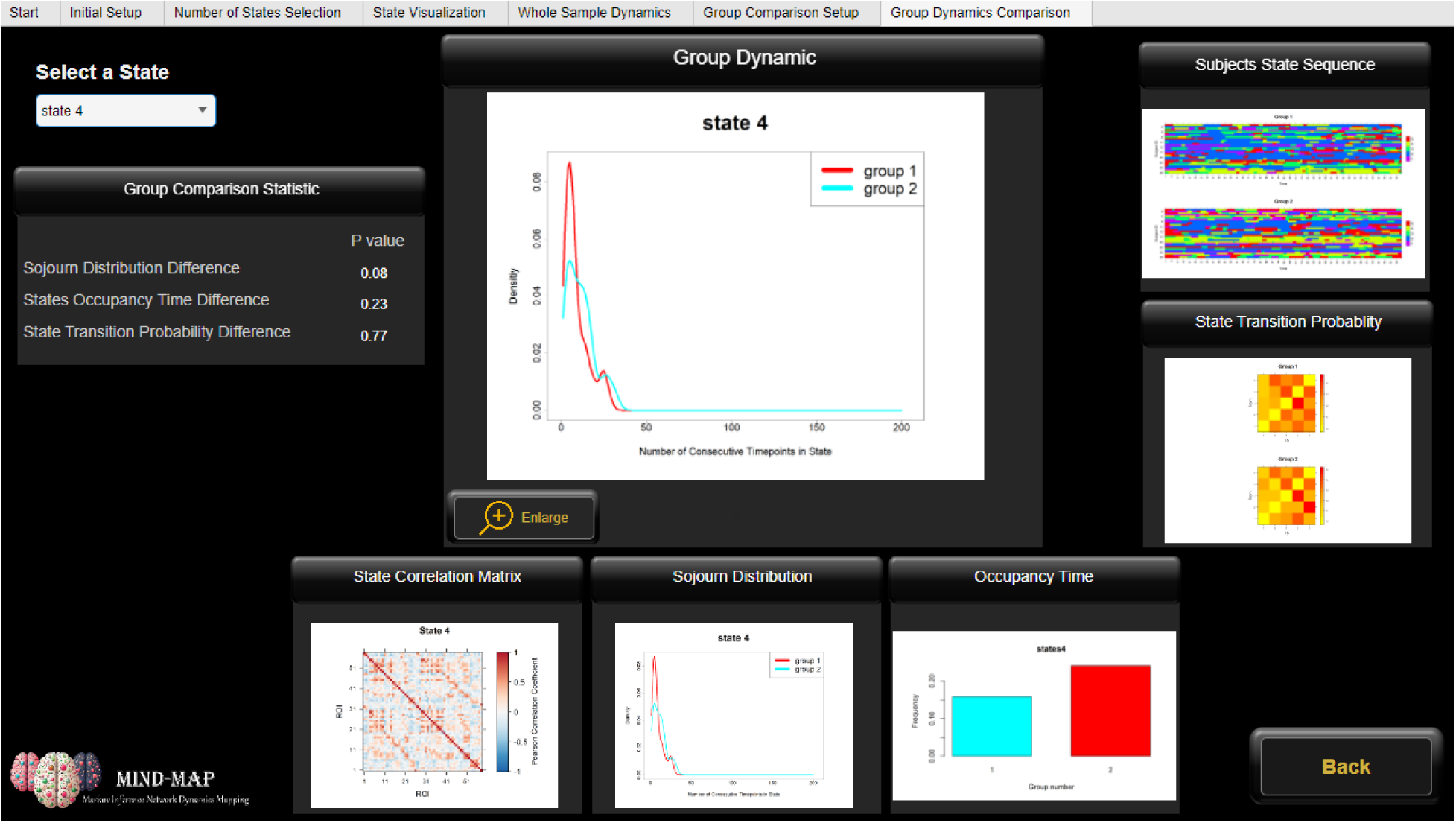
Two-sample dynamics comparison. This includes graphical comparison between the groups in addition to two group comparison statistics.

The subject occupancy time and the number of transitions are compared between the two groups using 10,000 permutation tests. For each group, the difference in average occupancy time and the number of transitions is calculated and tested against the average difference from the permutation samples to estimate their significance.

## 7 Toolbox runtime

The toolbox runtime depends on several parameters, including the number of subjects, regions, timepoints, states, the state initialization method, and the sojourn distribution model. However, in general, it runs within a reasonable amount of time. For example, running the HSMM model, using equal partitioning as the initial states estimation method, on a sample of 76 subjects (two paired groups of 38 subjects each), with 58 regions and 191 time points per region, using the toolbox’s default settings on a home desktop with an Intel(R) Core(TM) i5-8500T CPU @ 2.11 GHz, took approximately 51 minutes to complete for a range of 3 to 10 states (8 runs of the HSMM model with different numbers of states). Typically, running the model on the entire sample takes significantly longer than running it on subgroups. This is because, in subgroup analysis, the states are fixed, allowing the toolbox to focus solely on identifying the optimal dynamics for each group without estimating new states. Furthermore, it does not require testing different numbers of states, as the number of states is determined in a prior step. For the same sample and setup, the subgroup analysis, including two-sample statistical testing with permutation testing for sojourn distribution, state occupancy time, and the number of transitions, took approximately 9 minutes to complete with 6 states.

## 8 Discussion

The primary goal of developing new statistical and mathematical methods for fMRI data analysis is to make them publicly available for diverse neuroscience research in a user-friendly format. Although there are several toolboxes for the investigation of static brain networks using fMRI data, such as Conn [16], tools for analyzing dynamic brain states are very limited. To address this gap, we developed the MIND-Map toolbox, which dynamically presents the results for easy user exploration. This toolbox utilizes the HSMM and generates a wide range of outputs to explore different aspects of brain states, including state connectivity matrices, brain maps, topological network measures, and their dynamics such as subject state sequence, state transition probabilities, state occupancy times, and sojourn distributions. The toolbox also supports two-sample statistical testing, enabling comparisons between patient and control groups or pre- and post-condition dynamics. We also introduced and implemented a new approach for identifying the optimal number of states when HSMM is applied to the fMRI data, which addresses the challenge of selecting the hidden number of states.

In addition to toolbox development, we performed several simulation experiments to investigate the HSMM model performance, which has not been investigated. We used a novel methodology to generate a new set of fMRI simulated data that we believe is much more realistic to real data compared to simulated data used in previous dynamic connectivity studies. More Specifically, instead of states with random topology or modular structures, we embedded real static brain networks in the time series. We also used an estimated subject sequence from one of our studies on real fMRI data instead of random subject state sequences. These two characteristics made our simulated data unique. Using this data, we showed that HSMM can accurately estimate the hidden states in the presence of a random noise of up to 50%. In addition, we showed the importance of selecting the number of states when the HSMM model is used. Finally, we used this simulated data set to demonstrate our approach’s accuracy in finding the optimal number of states.

## 9 Limitations

Although the toolbox simplifies the implementation of the HSMM and visualization of the results and introduces a novel approach for identifying the optimal number of states, several limitations remain. First, the HSMM model requires a relatively large dataset when analyzing the entire brain. Therefore, when working with smaller samples (fewer than 100 participants), it is not feasible to include all brain regions in the analysis simultaneously using highly parcellated atlases. This limitation could be addressed by estimating a sparse version of the inverse covariance matrix for the states. Also, the current version of the toolbox only supports two-group comparisons, and future updates could expand its capabilities to include more than two groups. Finally, the toolbox can be further developed to analyze the association between states’ dynamics and covariates of interest.

In addition to the toolbox limitations, there are some limitations with our simulation analysis. While our approach for identifying the true number of states worked well for the two simulated datasets, it requires a sufficient sample size and distinct brain states, which may not always be the case. Additionally, while our simulated dataset includes unique characteristics that enhance its realism compared to other functional dynamic simulation studies, we used random Gaussian noise, as the true noise model in fMRI data is not yet well understood. Future studies may focus on exploring alternative noise models to improve simulation accuracy.

## Acknowledgments

This research was supported by NIH grants P50AA026117 and K25EB032903.

## Data Availability

The toolbox and the simulated datasets used in this study will be made available upon acceptance of the manuscript.

