## Supplementary Figure 1 for "MIND-Map; A Comprehensive Toolbox for Estimating Brain Dynamic States"

**Supplementary Material.**

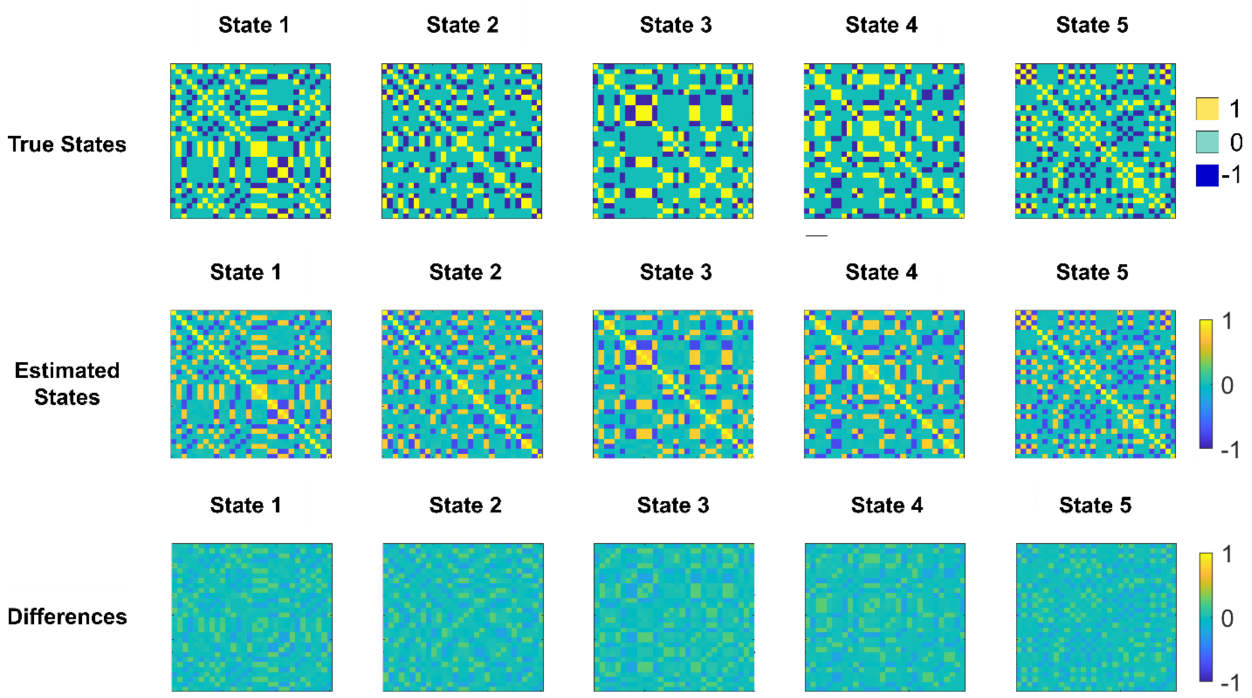


*Figure 1. Performance of the HSMM on a single set of the SIM-1 dataset which includes 10% percent random noise. Top. True states correlation matrices. Middle. The estimated states. Bottom the difference between the true states and estimated states*
