## Supplementary Figure 2 for "MIND-Map; A Comprehensive Toolbox for Estimating Brain Dynamic States"

**Supplementary Material.**

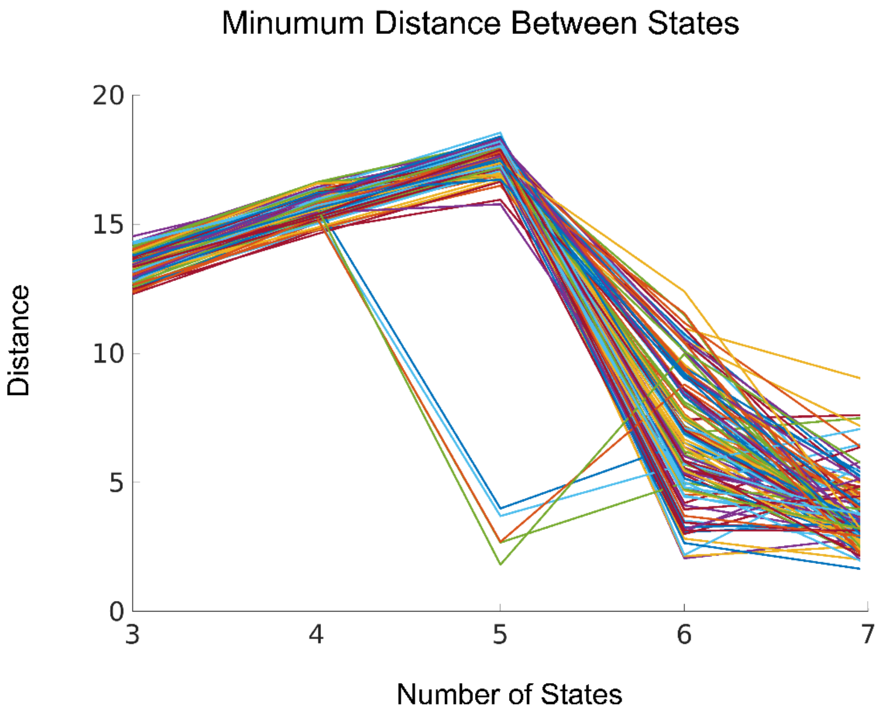


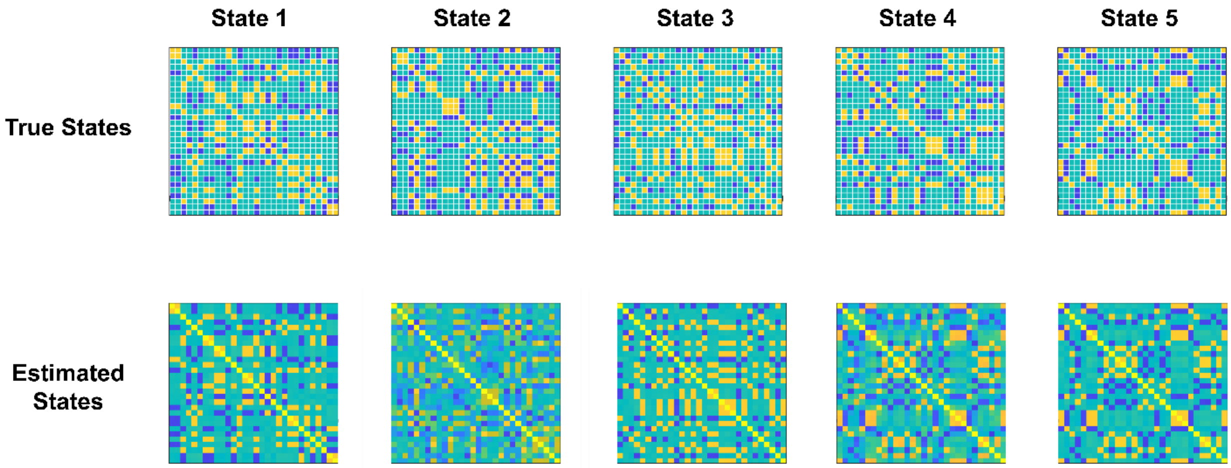


***Figure 2.*** *We initially generated 120 groups using the SIM-1 approach. However, due to the long runtime of various HSMM simulations, the main manuscript analysis focused on 50 groups for both SIM-1 and SIM-2.* ***Top:*** *When our method was applied to 120 groups from SIM-1 to identify the true number of states, it failed in 5 out of the 120 groups.* ***Bottom:*** *A closer examination of the estimated states revealed that these failures were caused by the HSMM model's inability to accurately estimate the true states, rather than a limitation of our method in estimating the optimal number of states.*
